# Expansion of the fatty acyl reductase gene family shaped pheromone communication in Hymenoptera

**DOI:** 10.1101/362509

**Authors:** Michal Tupec, Aleš Buček, Heiko Vogel, Václav Janoušek, Darina Prchalová, Jiří Kindl, Tereza Pavlíčková, Petra Wenzelová, Ullrich Jahn, Irena Valterová, Iva Pichová

## Abstract

The conserved fatty acyl reductase (FAR) family is involved in biosynthesis of fatty alcohols that serve a range of biological roles. In moths, butterflies (Lepidoptera), and bees (Hymenoptera), FARs biosynthesize fatty alcohol pheromones participating in mate-finding strategies. Using a combination of next-generation sequencing, analysis of transposable elements (TE) in the genomic environment of FAR genes, and functional characterization of FARs from *Bombus lucorum, B. lapidarius*, and *B. terrestris*, we uncovered a massive expansion of the FAR gene family in Hymenoptera, presumably facilitated by TEs. Expansion occurred in the common ancestor of bumblebees (Bombini) and stingless bees (Meliponini) after their divergence from the honeybee lineage. We found that FARs from the expanded FAR-A orthology group contributed to the species-specific male marking pheromone composition. Our results indicate that TE-mediated expansion and functional diversification of the FAR gene family played a key role in the evolution of pheromone communication in the crown group of Hymenoptera.

**Abbreviations:** MMP: male marking pheromone, FA: fatty acid, FAME: fatty acid methyl ester, FAR: fatty acyl reductase, LG: labial gland, FB: fat body, TE: transposable element.

## Introduction

Accumulation of DNA sequencing data is greatly outpacing our ability to experimentally assess the function of the sequenced genes, and most of these genes are expected to never be functionally characterized^1^. Important insights into the evolutionary processes shaping the genomes of individual species or lineages can be gathered from predictions of gene families, gene orthology groups and gene function. However, direct experimental evidence of the function of gene family members is often unavailable or limited^2–6^. Gene duplication is hypothesized to be among the major genetic mechanisms of evolution^7,8^. Although the most probable evolutionary fate of duplicated genes is the loss of one copy, the temporary redundancy accelerates gene sequence divergence and can result in gene subfunctionalization or neofunctionalization—acquisition of slightly different or completely novel functions in one copy of the gene^9,10^.

The alcohol-forming fatty acyl-CoA reductases (FARs, EC 1.2.1.84) belong to a multigene family that underwent a series of gene duplications and subsequent gene losses, pseudogenizations and possibly functional diversification of some of the maintained copies, following the birth-and-death model of gene family evolution^11^. FARs exhibit notable trends in gene numbers across organism lineages; there are very few FAR genes in fungi, vertebrates and non-insect invertebrates such as *Caenorhabditis elegans*, whereas plant and insect genomes typically harbour extensive FAR gene family members^11^. FARs are critical for production of primary fatty alcohols, which are naturally abundant fatty acid (FA) derivatives with a wide variety of biological roles. Fatty alcohols are precursors of waxes and other lipids that serve as surface-protective or hydrophobic coatings in plants, insects and other animals^12–14^; precursors of energy-storing waxes^15–17^; and components of ether lipids abundant in the cell membranes of cardiac, nervous and immunological tissues^18^.

Additionally, in some insect lineages, FARs were recruited for yet another task—biosynthesis of fatty alcohols that serve as pheromones or pheromone precursors. Moths (Lepidoptera) are the most well-studied model of insect pheromone biosynthesis and have been the subject of substantial research effort related to FARs. Variation in FAR enzymatic specificities is a source of sex pheromone signal diversity among moths in the genus *Ostrinia*^19^ and is also responsible for the distinct pheromone composition in two reproductively isolated races of the European corn borer *Ostrinia nubilalis*^20^. Divergence in pheromone biosynthesis can potentially install or strengthen reproductive barriers, ultimately leading to speciation^21^. However, the biological significance of a large number of insect FAR paralogs remains unclear, as all FARs implicated in moth and butterfly sex pheromone biosynthesis are restricted to a single clade, indicating that one FAR group was exclusively recruited for pheromone biosynthesis^20,22–24^. While more than 20 FARs have been experimentally characterized from 23 moth and butterfly (Lepidoptera) species^25^, FARs from other insect orders have received far less attention. Single FAR genes have been isolated and experimentally characterized from *Drosophila* (Diptera)^14^, the European honeybee (Hymenoptera) ^26^ and the scale insect *Ericeus pela* (Hemiptera)^27^. Our limited knowledge about FAR function prevents us from drawing inferences about the biological significance of the FAR gene family expansion in insects.

Bumblebees (Hymenoptera: Apidae) are a convenient experimental model to study insect FAR evolution because the majority of bumblebee species produces fatty alcohols as species-specific components of male marking pheromones (MMPs)^28^, which are presumed to be biosynthesized by some of the numerous bumblebee FAR gene family members^29^. Bumblebee males employ MMPs to attract conspecific virgin queens^30^. In addition to fatty alcohols, MMPs generally contain other FA derivatives and terpenoid compounds. The MMP composition serves as a phylogenetic signal and can be used as a taxonomic tool to discriminate bumblebee species and subspecies^31–33^. Pheromones from three common European bumblebee species, *B. terrestris, B. lucorum* and *B. lapidarius*, represent the diversity of fatty alcohol MMP components. Fatty alcohols are the major compounds in MMPs of *B. lapidarius* (hexadecanol and *Z*9-hexadecenol) and accompanying electroantennogram-active compounds in *B. terrestris* (hexadecanol, octadecatrienol, octadecenol) and *B. lucorum* (hexadecanol, *Z*9,*Z*12-octadecadienol, *Z*9,*Z*12,*Z*15-octadecatrienol, octadecanol)^34–40^.

In our previous investigation of the molecular basis of pheromone diversity in bumblebees, we found that the substrate specificities of fatty acyl desaturases (FADs), enzymes presumably acting upstream of FARs in pheromone biosynthesis^41^, are conserved across species despite differences in the compositions of their unsaturated FA-derived pheromone components^42^. These findings suggest that the substrate specificity of FADs expressed in the male bumblebee MMP-producing labial gland (LG) contributes only partially to the species-specific composition of FA-derived MMPs^42^. The fatty alcohol content in bumblebee MMPs is therefore presumably co-determined by the enzymatic specificity of other pheromone biosynthetic steps, such as FA biosynthesis/transport or FA reduction. Analysis of the *B. terrestris* male LG transcriptome uncovered a remarkably high number of putative FAR paralogs, including apparently expressed pseudogenes, strongly indicating dynamic evolution of the FAR gene family^29^.

Here, we aimed to determine how the members of the large FAR gene family in the bumblebee lineage contribute to MMP biosynthesis. We sequenced and analysed *B. lucorum* and *B. lapidarius* male LG and FB transcriptomes and functionally characterized the FAR enzymes, along with FAR candidates from *B. terrestris*, in a yeast expression system. We combined experimental information about FAR enzymatic specificities with quantitative information about bumblebee FAR expression patterns, as well as comprehensive GC analysis of MMPs and their FA precursors in the bumblebee male LG, with inference of the hymenopteran FAR gene tree. In addition, we investigated the content of transposable elements (TE) in the genomic environment of FAR genes in *B. terrestris*. We concluded that a dramatic TE-mediated expansion of the FAR gene family started in the common ancestor of the bumblebee (Bombini: *Bombus*) and stingless bee (Meliponini) lineages, which presumably shaped the pheromone communication in these lineages.

## Results

### Identification of FARs in bumblebee transcriptomes

We sequenced, assembled and annotated male LG and male fat body (FB) transcriptomes of two bumblebee species, *B. lucorum* and *B. lapidarius*. LG is the MMP-producing organ and is markedly enlarged in males, while FB was used as a reference tissue not directly involved in MMP biosynthesis^43^. Searches of the LG and FB transcriptomes of *B. lucorum* and *B. lapidarius* and the previously sequenced FB and LG transcriptomes of *B. terrestris*^29^ yielded 12, 26, and 16 expressed FAR homologs in *B. lapidarius, B. terrestris*, and *B. lucorum*, respectively (Supplementary Fig. 1).

### FAR gene family evolution in Hymenoptera

To gain insight into the evolution of FAR gene family in Hymenoptera, we reconstructed a FAR gene tree using predicted FARs from species representing ants, wasps, parasitic wasps and several bee lineages (Fig. 1). We assigned the names FAR-A to FAR-K to 11 FAR orthology groups that were retrieved as branches with high bootstrap support in the FAR gene tree. These orthology groups typically encompass one or more orthologs from each of the hymenopteran species used in the tree inference, with the exception of apparent species-specific FAR duplications or losses (Fig. 1). Notably, we identified a massive expansion of the FAR-A orthology group in the bumblebee and stingless bee (subfamily Meliponini) lineages, the two most closely related lineages in our dataset. The number of FAR homologs is inflated by a large number of FAR transcripts with incomplete protein coding sequences lacking catalytically critical regions such as the putative active site, NAD(P)^+^ binding site or substrate binding site (Fig. 1, Supplementary Table 1). We also inferred a FAR gene tree encompassing FARs from three representatives of non-hymenopteran insect orders—the beetle *Tribolium castaneum*, the moth *Bombyx mori* and the fly *Drosophila melanogaster* (Supplementary Fig. 2). The only functionally characterized FAR from *D. melanogaster*—Waterproof (NP_651652.2), which is involved in biosynthesis of a protective wax layer^14^—was placed in the FAR-J orthology group (Supplementary Fig. 2). The FAR-G orthology group includes a FAR gene from *Apis mellifera* with unclear biological function^26^ and a sex pheromone-biosynthetic FAR from *B. mori*^44^ (Supplementary Fig. 2). In the gene tree, the majority of FAR orthology groups contain predicted FARs from both hymenopteran and non-hymenopteran insect species. These orthology groups are presumably ancestral to insects. Only FAR-D and FAR-K do not include any non-hymenopteran FARs from our dataset and thus presumably represent hymenoptera-specific FAR gene family expansions (Supplementary Fig. 2).

**Figure 1.**
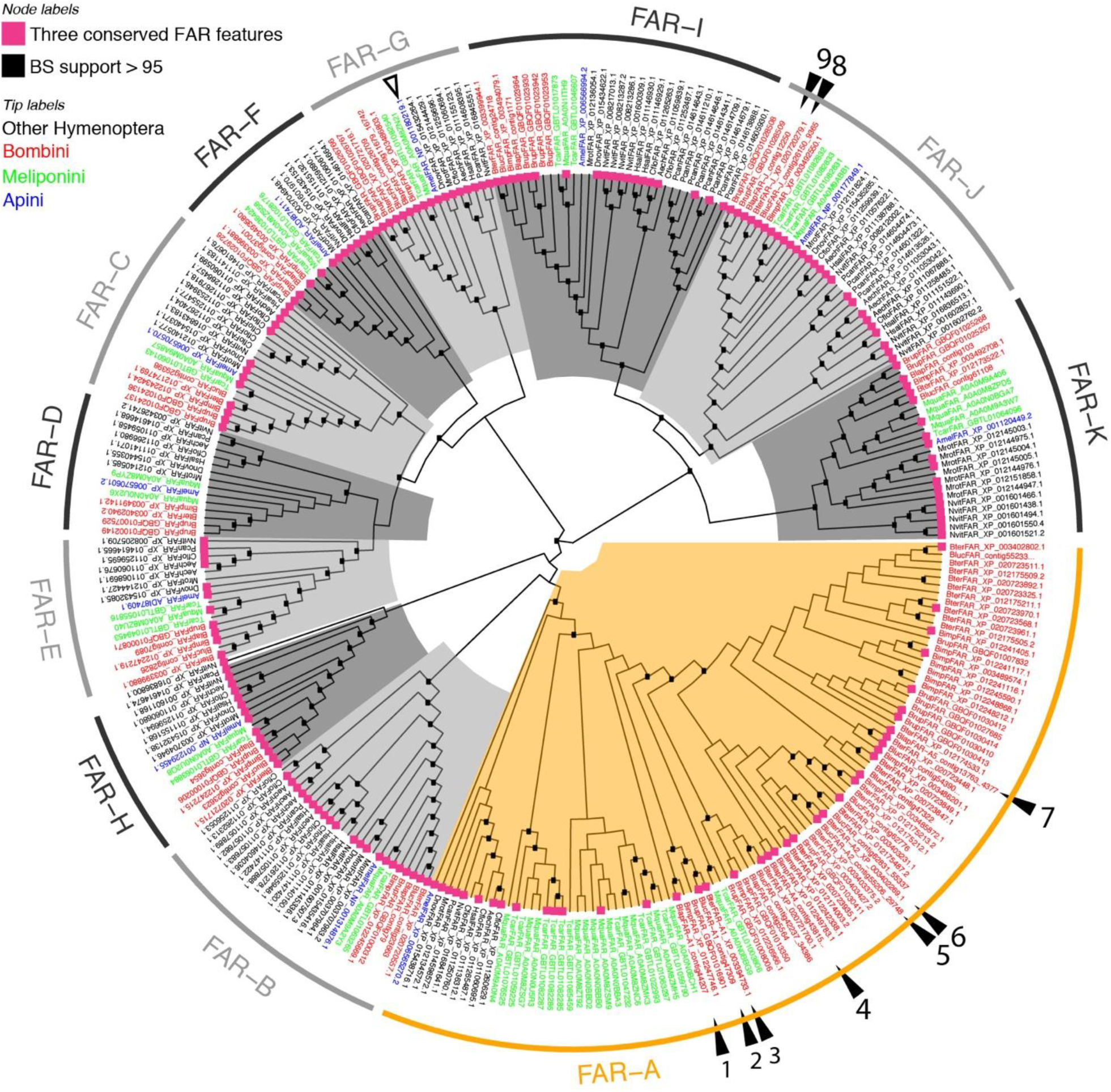
Hymenopteran FAR gene tree. Tree tips are coloured according to taxonomy: red, bumblebee FARs (*B. terrestris, B. lucorum, B. lapidarius, B. impatiens, B. rupestris*); green, stingless bee FARs (*Tetragonula carbonaria, Melipona quadrifasciata*); blue, *A. mellifera* FARs; and black, FARs from other hymenopteran species. The FAR-A orthology group is highlighted orange; other orthology groups in shades of grey. Functionally characterized bumblebee FARs from this study are indicated by filled triangles and numbered.1: *Blap*FAR-A1, 2: *Bluc*FAR-A1, 3: *Bter*FAR-A1, 4: *Blap*FAR-A4, 5: *Bluc*FAR-A2, 6: *Bter*FAR-A2, 7: *Blap*FAR-A5, 8: *Bter*FAR-J, and 9: *Blap*FAR-J. The functionally characterized *A. mellifera* FAR is indicated by an empty triangle. Internal nodes highlighted with black boxes indicate bootstrap support >95%. Violet squares at the tree tips indicate FARs for which CDD search yielded all three FAR conserved features—active site, putative NAD(P)^+^ binding site and substrate binding site (see Table S1 for complete CDD search results).

### Genomic organization and TE content

To uncover the details of genetic organization of FAR-A genes, we attempted to analyse the shared synteny of FAR genes in the genomes of *B. terrestris* and *A. mellifera*^45^. We aligned the *A. melifera* and *B. terrestris* genomes, but we were not able to identify any positional *A. mellifera* homologs of *B. terrestris* FAR-A genes (data not shown). While the majority of FAR genes belonging to the non-FAR-A gene orthology group localize to the *B. terrestris* genome assembled to linkage groups, most of the *B. terrestris* FAR-A genes localize to unlinked short scaffolds (Supplementary Table 2). Some of the FAR-A genes in the *B. terrestris* genome are arranged in clusters (Supplementary Fig. 3).

A genome assembly consisting of short scaffolds is often indicative of a repetitive structure in the assembled genomic region. Our analysis of the distribution of TEs in the vicinity of FAR genes in the *B. terrestris* genome confirmed that TEs are significantly enriched around FAR-A genes compared to the genome-wide average around randomly selected genes (*p* < 0.0001). FAR-A genes have on average more than 50% of their 10-kb surrounding region formed by TEs compared to an average 10% around randomly selected *B. terrestris* genes. In contrast, the densities of TEs in the vicinity of FAR genes not belonging to the FAR-A group do not differ from the genome-wide average (*p* = 0.1041; Fig. 2). Although all major known TE families are statistically enriched in the neighbourhood of the FAR-A genes (Fig. 2), the Class I comprising retroid elements contributes considerably to the elevated repeat content around FAR-A genes.

**Figure 2.**
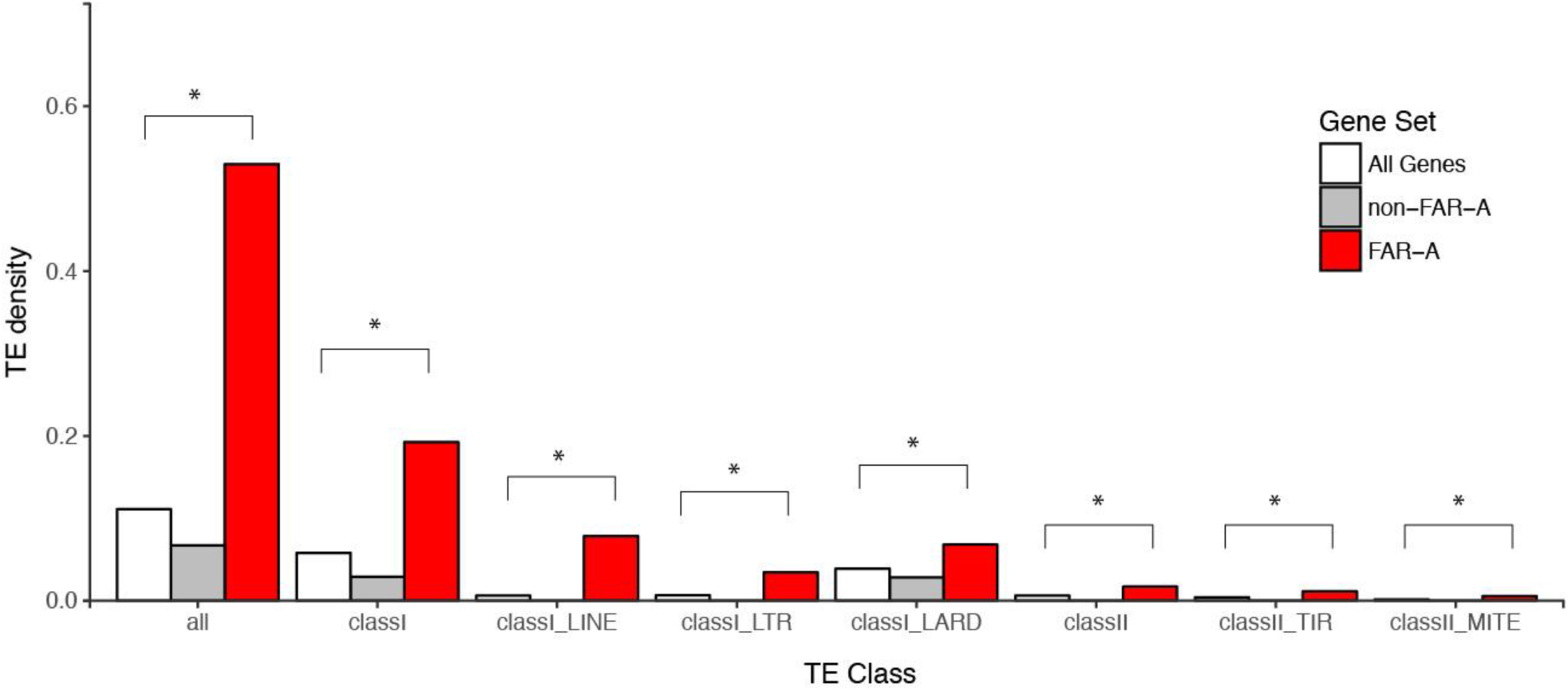
Average TE densities in 10 kb windows around groups of *B. terrestris* genes. “All Genes” represents randomly selected sets of *B. terrestris* genes, and non-FAR-A includes FAR genes belonging to the non-FAR-A orthology groups. Densities were analyzed for all TEs (all) and separately for Class I, Class II and the most abundant TE families within each class (LINEs, LTRs, LARDs, TIRs, MITEs). Significant differences (*p* < 0.05, two-tailed *t*-test) are marked with asterisks.

### Tissue-specific expression

We selected 10 promising MMP-biosynthetic FAR candidates that were 1) among the 100 most abundant transcripts in the LG and were substantially more abundant in LG than in FB based on RNA-Seq-derived normalized expression values (Supplementary Fig. 1 and ref. ^29^) and 2) included in the protein coding sequence all the predicted catalytically critical regions of FARs—the putative active site, NAD(P)^+^ binding site and substrate binding site (Supplementary Table 1).

By employing RT-qPCR in an expanded set of bumblebee tissues, we confirmed that the FAR candidates follow a general trend of overexpression in male LG compared to FB, flight muscle and gut (all from male bumblebees) and virgin queen LG (Fig. 3, Supplementary Fig. 4, *p* < 0.05, one-way ANOVA followed by *post-hoc* Tukey’s HSD test). Notably, *B. lapidarius* FAR-A1 (*Blap*FAR-A1) and *B. terrestris* FAR-J (*Bter*FAR-J) transcripts are also abundant in virgin queen LG, where they are expressed at levels comparable to those in male LG (Fig. 3, Supplementary Fig. 4).

**Figure 3.**
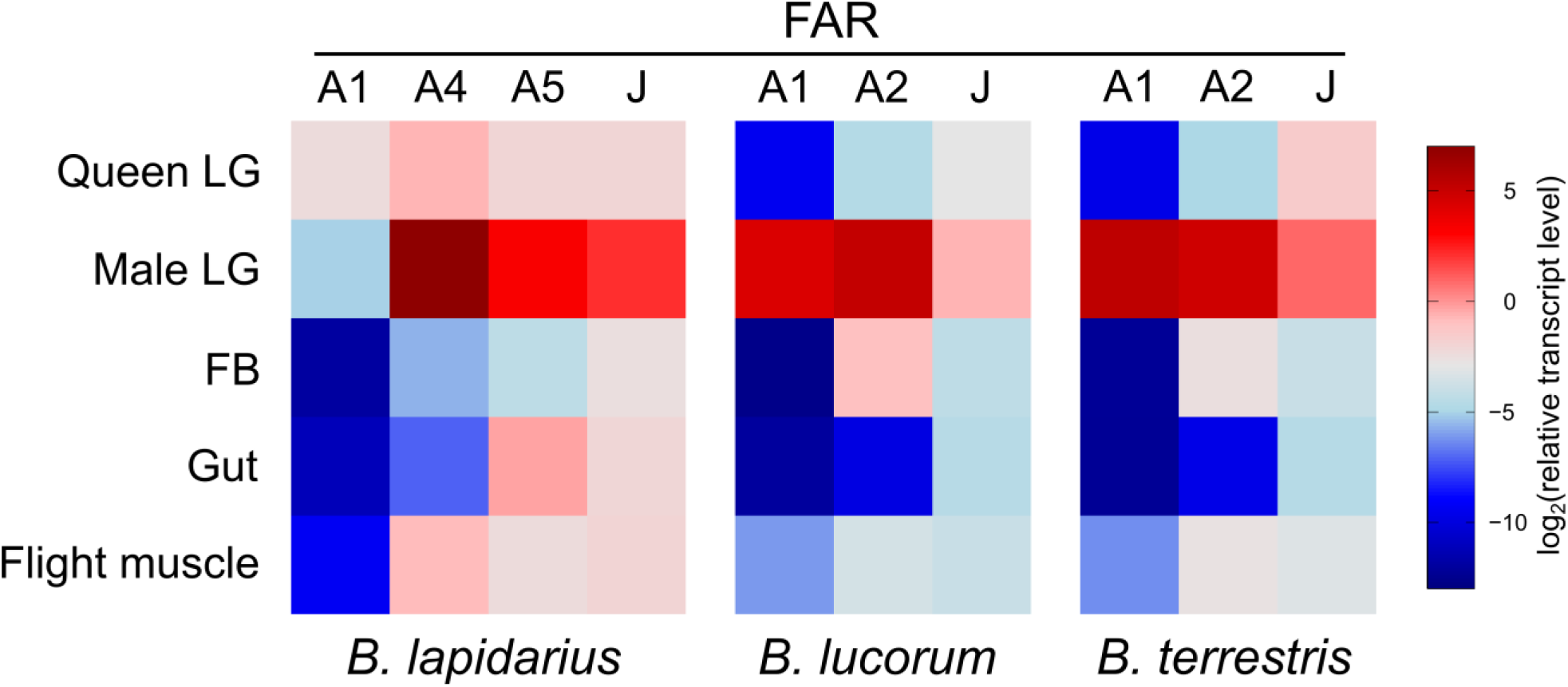
Relative transcript levels of FAR candidates across bumblebee tissues. The transcript levels were assayed by RT-qPCR in male tissues (LG, FB, gut, flight muscle) and in LGs of virgin queens.

### Cloning and functional characterization

The full-length coding regions of the FAR candidates were isolated from male LG cDNA libraries using gene-specific PCR primers (Supplementary Table 3). In general, the FAR candidates share high to very high protein sequence similarity within each orthology group. FARs from three bumblebee species belonging to the FAR-J orthology group are nearly identical, sharing 97.2–99.7% protein sequence identity; *Bluc*FAR-A1 and *Bter*FAR-A1 share 99.4% protein sequence identity with each other and 60.9–61.1% with *Blap*FAR-A1. *Bluc*FAR-A2 and *Bter*FAR-A2 share 94.8% protein sequence identity (Supplementary Table 4). *Bluc*FAR-J was not cloned because of its very high similarity to *Bter*FAR-J (99.7% sequence identity, two amino acid differences). We cloned two versions of *Blap*FAR-A1: one that was custom-synthesized based on the predicted full-length coding sequence assembled from RNA-Seq data and one called *Blap*FAR-A1-short that we consistently PCR-amplified from *B. lapidarius* male LG cDNA. *Blap*FAR-A1-short has an in-frame internal 66 bp deletion in the coding region that does not disrupt the predicted active site, putative NAD(P)^+^ binding site or putative substrate binding site. Using RT-qPCR with specific primers for each variant, we confirmed that both *Blap*FAR-A1 and *Blap*FAR-A1-short are expressed in the *B. lapidarius* male LG and virgin queen LG (Supplementary Fig. 5).

To test whether the MMP-biosynthetic FAR candidates code for enzymes with fatty acyl reductase activity and to uncover their substrate specificities, we cloned the candidate FAR coding regions into yeast expression plasmids, heterologously expressed the FARs in *Saccharomyces cerevisiae* and assayed the fatty alcohol production by GC (Supplementary Fig. 8 and Supplementary Fig. 9). His-tagged FARs were detected in all yeast strains transformed with plasmids bearing FARs (Fig. 4, Supplementary Fig. 11a), while no His-tagged proteins were detected in the negative control (yeast strain transformed with an empty plasmid). In addition to the major protein bands corresponding to the theoretical FAR molecular weight, we typically observed protein bands with lower and/or higher molecular weight (Fig. 4a). The synthetic *Bluc*FAR-A1-opt and *Bluc*FAR-A2-opt coding regions with codon usage optimized for *S. cerevisiae* showed a single major Western blot signal corresponding to the position of the predicted full-length protein (Fig. 4a). The shortened heterologously expressed proteins thus presumably represent incompletely transcribed versions of full-length FARs resulting from ribosome stalling^46^, while the higher molecular weight bands might correspond to aggregates of full-length and incompletely translated FARs. Because the codon-optimized *Bluc*FAR-A1-opt and *Bluc*FAR-A2-opt exhibit the same overall specificity in yeast expression system as the respective non-codon-optimized FARs (Supplementary Fig. 6), we employed non-codon-optimized FARs for further functional characterization. We only used the codon-optimized versions of *Bluc*FAR-A1 and *Bluc*FAR-A2 in experiments with exogenously supplemented substrates to increase the possibility of product detection, as the optimized FARs produce overall higher quantities of fatty alcohols (*p* < 0.05, two-tailed *t*-test).

**Figure 4.**
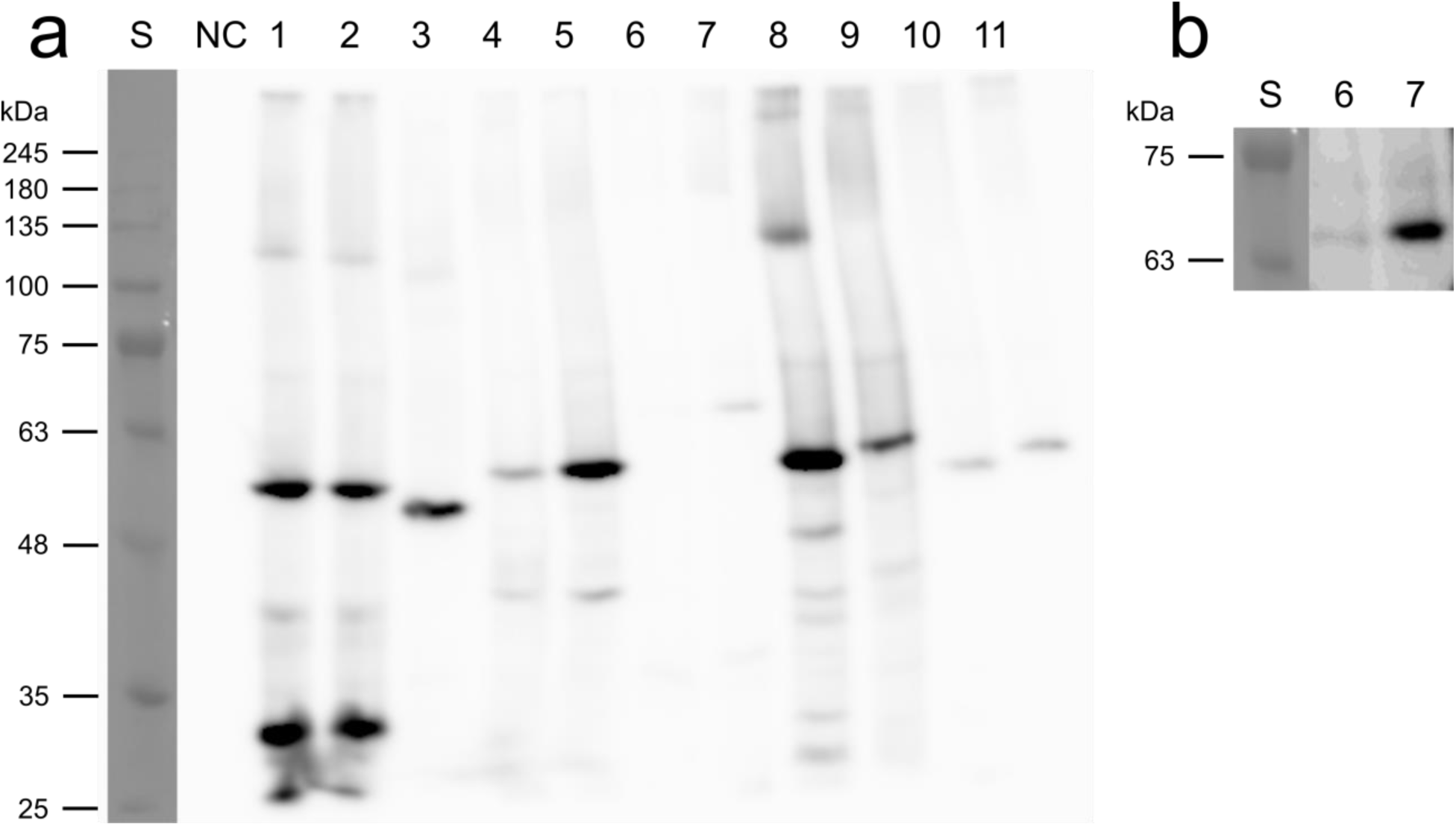
Western blot analysis of FAR protein expression in yeast cell lysates. (**a**) Photograph of the membrane after detection with anti-6×His-tag antibody. (**b**) Detail of the FAR-J full-length protein region with increased contrast. *Lanes*: S, protein standard (VI, AppliChem); NC, negative control (yeast carrying empty vector); 1–11, yeast strains carrying plasmids with *Bluc*FAR-A1 (1), *Bter*FAR-A1 (2), *Blap*FAR-A1 (3), *Bluc*FAR-A2 (4), *Bter*FAR-A2 (5), *Bter*FAR-J (6), *Blap*FAR-J (7), *Blap*FAR-A4 (8), *Blap*FAR-A5 (9), *Bluc*FAR-A1-opt (10) and *Bluc*FAR-A2-opt (11).

Characterization of FAR enzymatic activities involved identification of numerous individual FA derivatives, denoted using the position/configuration of the double bond if present (e.g., *Z*9), the length of the carbon chain (e.g., 20), the number of double bonds (e.g., “:” or “:1” for saturated and monounsaturated FAs, respectively) and the C1 moiety (COOH for acid, OH for alcohol, Me for methyl ester, CoA for CoA-thioester).

Functional characterization of FARs from *B. terrestris* and *B. lucorum* in yeast indicated that saturated C16 to C26 fatty alcohols are produced by both *Bter*/*Bluc*FAR-A1 and *Bter*FAR-J enzymes (Fig. 5ad, Supplementary Fig. 9); *Bter*/*Bluc*FAR1 prefers C22 substrates, whereas *Bter*FAR-J has an optimal substrate preference slightly shifted to C24. Unlike any of the other characterized FARs, *Bter*/*Bluc*FAR-A1 are also capable of reducing supplemented monounsaturated *Z*15-20:1 acyl to the corresponding alcohol (Fig. 5a and Supplementary Fig. 10c). Both *Bter*FAR-A2 and *Bluc*FAR-A2 reduce only 16: and 18: acyls (Fig. 5c, Supplementary Fig. 8).

**Figure 5.**
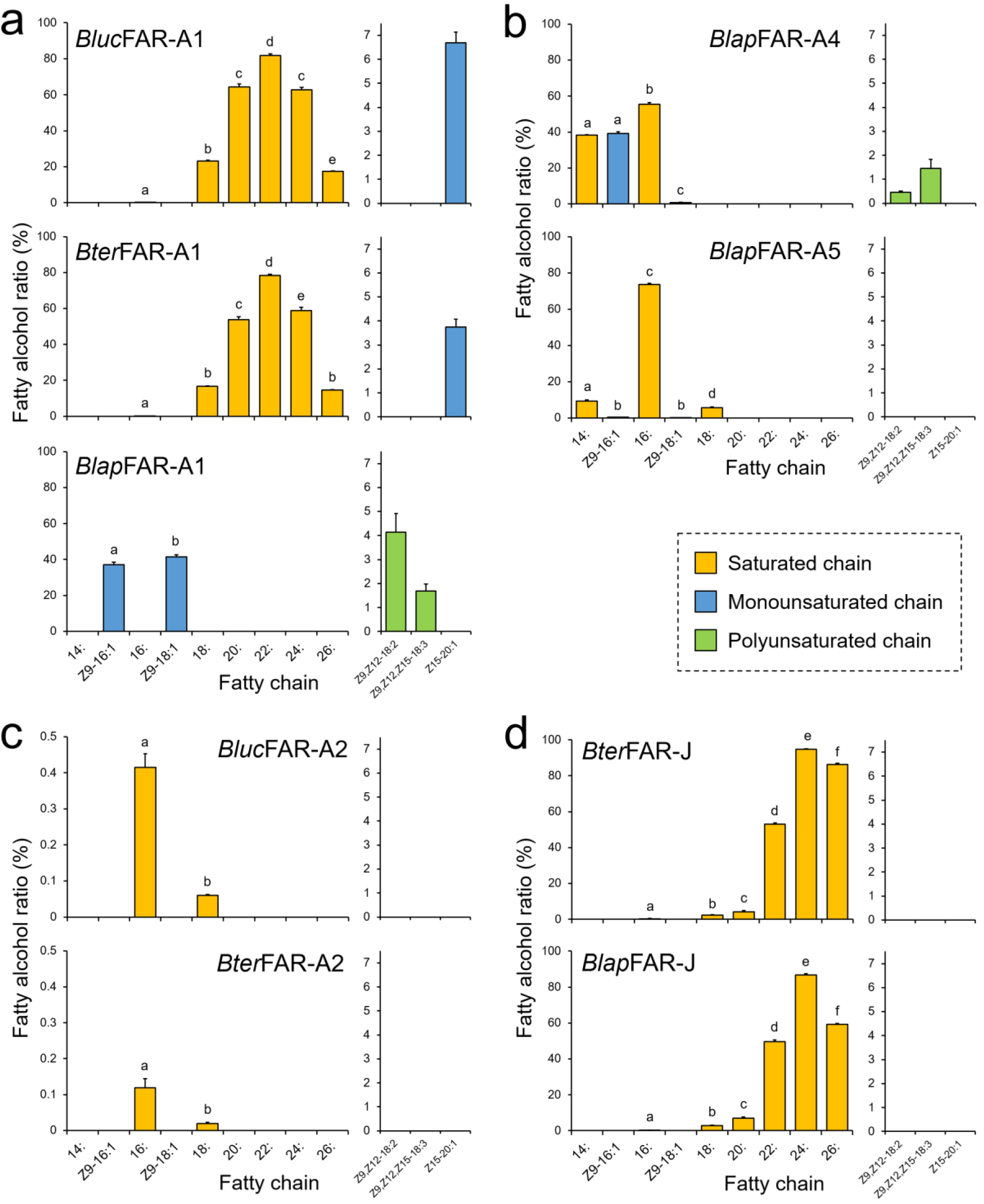
Fatty alcohol ratios in yeast strains expressing bumblebee FARs assayed on yeast native lipids and after supplementation of yeast with either *Z*9,*Z*12-18:2, *Z*9,*Z*12,*Z*15-18:3 or *Z*15-20:1 acyls. Significant differences (*p* < 0.01, one-way ANOVA followed by *post*-*hoc* Tukey’s HSD test) are marked with different letters. See Methods for description of fatty alcohol ratio calculation.

Characterization of *B. lapidarius* FARs showed that *Blap*FAR-A1, in contrast to *Bter*/*Bluc*FAR-A1, produces *Z*9-16:1OH and *Z*9-18:1OH (Fig. 5A and Supplementary Fig. 8). *Blap*FAR-A4 produces 16:OH and *Z*9-16:1OH, together with lower quantities of 14:OH and *Z*9-18:1OH (Fig. 5b, Supplementary Fig. 7, Supplementary Fig. 8). *Blap*FAR-A5 produces 16:OH as a major product and lower amounts of 14:OH, *Z*9-16:1OH, 18:OH and Z9-18:1OH (Fig. 5b, Supplementary Fig. 7, Supplementary Fig. 8). In addition, both *Blap*FAR-A1 and *Blap*FAR-A4 are capable of reducing supplemented polyunsaturated fatty acyls (*Z*9,*Z*12-18:2 and *Z*9,*Z*12,*Z*15-18:3) to their respective alcohols (Fig. 5ab and Supplementary Fig. 10ab). Similarly to *Bter*FAR-J, *Blap*FAR-J also reduces saturated C16 to C26 acyls (Fig. 5d and Supplementary Fig. 9). No fatty alcohols were detected in the negative control (Supplementary Fig. 8). We did not detect the formation of fatty aldehydes in any of the yeast cultures (data not shown), confirming that the studied FARs are strictly alcohol-forming fatty acyl-CoA reductases. In contrast to *Blap*FAR-A1, *Blap*FAR-A1-short does not produce detectable amounts of any fatty alcohol (Supplementary Fig. 11b), suggesting that the missing 22-amino acid region is crucial for the retention of FAR activity (Supplementary Fig. 8).

### Quantification of fatty alcohols and fatty acyls in bumblebee male LG and FB

In addition to functional characterization of FARs in a heterologous host, we performed a detailed analysis of transesterifiable fatty acyls (free FAs and fatty acyls bound in esters) and fatty alcohols in LGs and FBs of 3-day-old *B. lapidarius, B. terrestris* and *B. lucorum* males (Fig. 6, Supplementary Table 5 and Supplementary Table 6).

**Figure 6.**
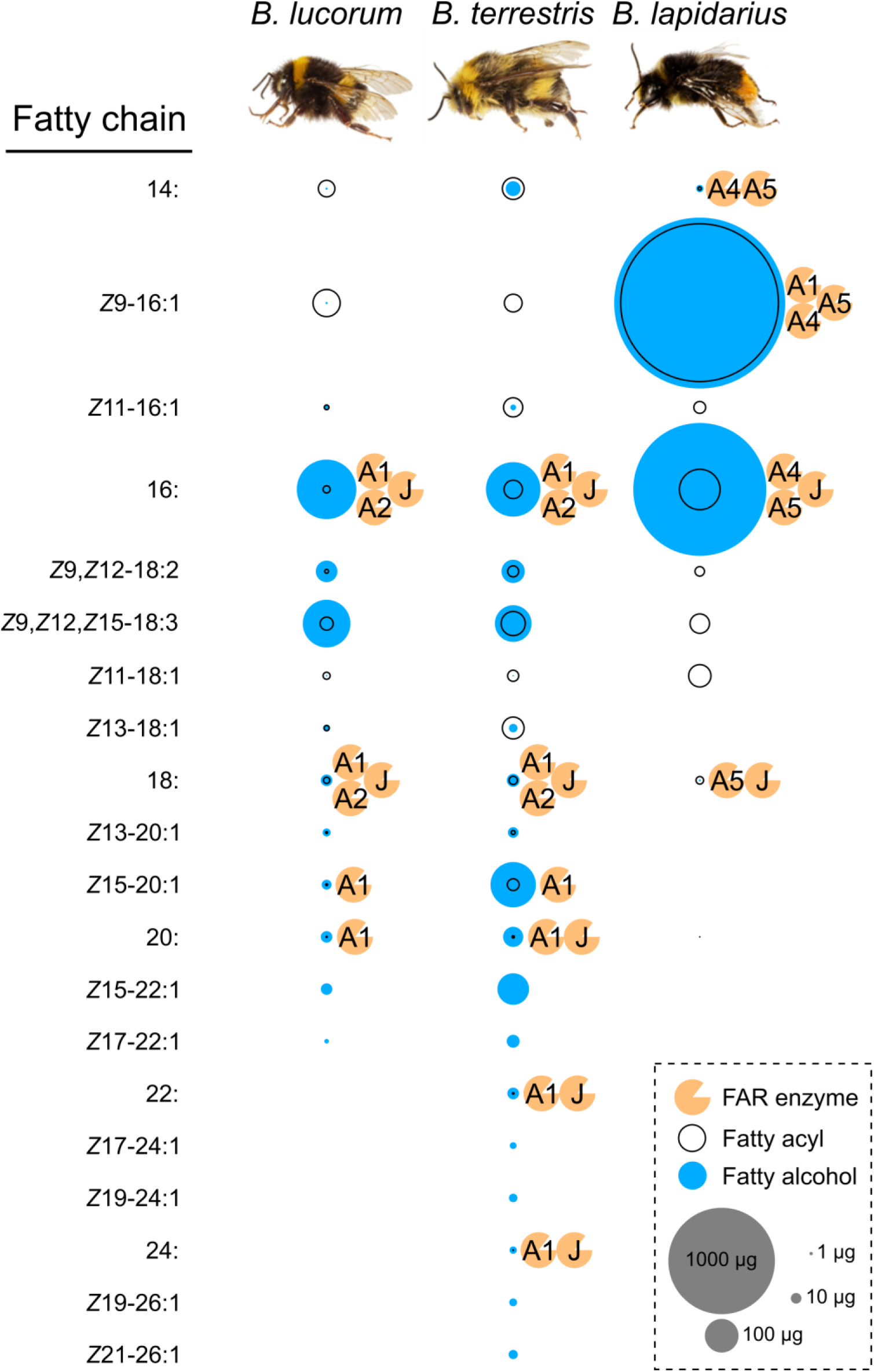
Fatty alcohol and fatty acyl composition in male LGs of *B. lucorum, B. terrestris* and *B. lapidarius* together with the proposed participation of FARs. The fatty acyls were determined as corresponding methyl esters. The size of fatty acyl and fatty alcohol circle, respectively, represents the average quantity in single male LG (Supplementary Table 5).

A limited number of fatty alcohols (mainly 16:OH, *Z*9,*Z*12-18:2OH and *Z*9,*Z*12,*Z*15-18:3OH) were detected in FBs of *B. lucorum* and *B. terrestris* (Supplementary Table 6), but at substantially lower abundance than in LGs. In the LGs, 4, 14, and 19 individual fatty alcohol compounds were detected in *B. lapidarius, B. lucorum* and *B. terrestris*, respectively (Supplementary Table 5). To assess the apparent *in-vivo* specificity of FARs in LGs and FBs, we calculated the ratios of amounts of each fatty alcohol to the amount of its hypothetical fatty acyl precursors (Supplementary Fig. 12). The fatty alcohol ratios are greater than 50% for most of the fatty alcohols in LGs and even approach 100% for some of the monounsaturated C20+ fatty alcohols (Supplementary Fig. 12ab), as the corresponding fatty acyls could not be quantified due to low abundance, suggesting that the FARs acting on these fatty chains convert almost all of the acyl substrate to alcohol. The specificities of the characterized FARs determined in yeast correlate well with the composition of LG fatty alcohols and fatty acyls (Fig. 6), except for *Z*9,*Z*12-18:2OH and *Z*9,*Z*12,*Z*15-18:3OH, as none of the studied FARs from *B. lucorum* or *B. terrestris* reduce the corresponding acyls.

## Discussion

Since the first genome-scale surveys of gene families, gene duplications and lineage-specific gene family expansions have been considered major mechanisms of diversification and adaptation in eukaryotes^2^ and prokaryotes^47^. Tracing the evolution of gene families and correlating them with the evolution of phenotypic traits has been facilitated by the growing number of next-generation genomes and transcriptomes from organisms spanning the entire tree of life. However, obtaining experimental evidence of the function of numerous gene family members across multiple species or lineages is laborious. Thus, such data are scarce, and researchers have mostly relied on computational inference of gene function^48^. Here, we aimed to combine computational inference with experimental characterization of gene function to understand the evolution of the FAR family, for which we predicted a notable gene number expansion in our initial transcriptome analysis of the buff-tailed bumblebee *B. terrestris*^29^. We specifically sought to determine whether the FARs that emerged through expansion of the FAR gene family substantially contribute to MMP biosynthesis in bumblebees.

We identified a massive expansion of the FAR-A orthologous gene group in stingless bees and bumblebees. The sister taxonomic relationship of these taxa and their position as a crown group within the bee (Anthophila) clade^49–51^ indicates that the FAR duplication process occurred or started in the common ancestor of bumblebees and stingless bees. According to estimated lineage divergence times, FAR duplication events started 76–85 million years ago after their divergence from *Apis* 82–93 million years ago^52^. The number of inferred FAR-A orthologs is inflated by predicted pseudogenes—FARs with fragmented coding sequences that lack some of the catalytically essential domains and motifs (Supplementary Fig. 2). These predicted catalytically inactive yet highly expressed FAR-A pseudogenes might play a role in regulating the FAR-catalysed reduction^53^. The number of predicted FAR-A pseudogenes indicate that the FAR-A orthology group expansion in this lineage was a highly dynamic process (Fig. 1, Supplementary Fig. 2). The high number of species-specific FAR-A duplications or losses between the closely related species *B. lucorum* and *B. terrestris*, which diverged approximately 5 million years ago^54^, further indicates the dynamic evolutionary processes acting on the FAR gene.

Strikingly, stingless bees also employ LG secretion in scent marking^55^. In worker stingless bees, LG secretion is used as a trail pheromone to recruit nestmates to food resources and generally contains fatty alcohols such as hexanol, octanol, and decanol in the form of their fatty acyl esters^56,57^. The correlation between FAR-A gene orthologous group expansion and use of LG-produced fatty alcohols as marking pheromones suggests a critical role for FAR-A gene group expansion in the evolution of scent marking. In the future, identification and characterization of FAR candidates involved in production of stingless bee worker LG-secretion could corroborate this hypothesis.

Bumblebee orthologs (FAR-G) of *B. mori* pheromone-biosynthetic FARs^44^ are not abundantly or specifically expressed in male bumblebee LGs, as evidenced by RNA-Seq RPKM values (Supplementary Fig. 1). MMP-biosynthetic FARs in bumblebees and female sex pheromone-biosynthetic FARs in moths (Lepidoptera) were most likely recruited independently for the tasks of pheromone biosynthesis.

Various models have attempted to describe the evolutionary mechanisms leading to the emergence and maintenance of gene duplicates^10^. The fragmented state of the *B. terrestris* genome and its limited synteny with the *A. mellifera* genome restricts our ability to reconstruct the genetic events accompanying the FAR duplications resulting in the FAR-A orthology group expansion. Taking into consideration the large quantities of MMPs in bumblebee males, we speculate that gene dosage benefits could substantially contribute to the duplications and duplicate fixation of MMP-biosynthetic FARs. Under this model, selection for increased amounts of fatty alcohols could fix the duplicated FARs in a population^2,10^. The MMP quantities in bees are substantially higher than quantities of pheromones in other insects of comparable size. For example, *B. terrestris, B. lucorum*, and *B. lapidarius* bumblebee males can produce several milligrams of MMPs (Kindl, personal communication), while the sphingid moth *Manduca sexta* produces tens of nanograms of sex pheromone^58^. Sexual selection favouring bumblebee males capable of producing large quantities of MMPs thus might have served as the evolutionary driver for repeated FAR-A duplication.

Mechanistically, gene duplications can be facilitated by associated TEs^59–61^. The content of repetitive DNA in the *B. terrestris* and *B. impatiens* genome assemblies is 14.8% and 17.9%, respectively ^62^, which is lower than in other insects such as the beetle *Tribolium castaneum* (30%), *Drosophila* (more than 20%) or the wasp *Nasonia vitripenis* (more than 30%) but substantially higher than in the honeybee *Apis mellifera* (9.5%)^63,64^. Our finding that TEs are enriched in the vicinity of FAR-A genes in the *B. terrestris* genome indicates that TEs presumably contributed to the massive expansion of the FAR-A orthology group (Fig. 2). One possible scenario is that the FAR-A gene in the common ancestor of bumblebees and stingless bees translocated to a TE-rich region, which subsequently facilitated expansions of this orthology group.

We have previously shown that the transcript levels of biosynthetic genes generally reflect the biosynthetic pathways most active in bumblebee LG^29,42^. For further experimental characterization, we therefore selected the FAR-A and FAR-J gene candidates, which exhibited high and preferential expression in male LG. The abundant expression of *Blap*FAR-A1 and *Bter*FAR-J in both virgin queen and male LG suggests that these FARs might also have been recruited for production of queen-specific signals^65^.

The spectrum of fatty alcohols in *B. terrestris* and *B. lucorum* male LG differs substantially from that of *B. lapidarius*. In both *B. terrestris* and *B. lucorum*, the male LG extract contains a rich blend of C14–C26 fatty alcohols with zero to three double bonds (Fig. 6, Supplementary Table 5). In *B. lapidarius*, the male LG extract is less diverse and dominated by *Z*9-16:1OH and 16:OH (Fig. 6 and Supplementary Table 5). The functional characterization of LG-expressed FARs uncovered how the distinct repertoire of LG-expressed FAR orthologs, together with differences in FAR substrate specificities, contributes to the biosynthesis of species-specific MMPs. We found that the highly similar *Bluc*/*Bter*FAR-A1 and *Blap*FAR-A1 orthologs exhibit distinct substrate preferences for longer fatty acyl chains (C18-C26) and shorter monounsaturated fatty acyl chains (*Z*9-16:1 and *Z*9-18:1), respectively. This substrate preference correlates with the abundance of *Z*9-16:1OH in *B. lapidarius* MMP and the almost complete absence of *Z*9-16:1OH in *B. lucorum* and *B. terrestris* (Supplementary Table 5). *Blap*FAR-A4 and to some extent *Blap*FAR-A5 likely further contribute to the biosynthesis of *Z*9-16:1OH in *B. lapidarius*. The ability of *Bluc*/*Bter*FAR-A1 (and not of *Blap*FAR-A1) to reduce long monounsaturated fatty acyls (*Z*15-20:1) also correlates with the absence of detectable amounts of *Z*15-20:1OH in *B. lapidarius* MMP.

Our comprehensive GC analysis of bumblebee male LGs, however, indicates that the composition of LG fatty acyls is another factor that contributes substantially to the final MMP composition. For example, the very low quantities of *Z*9-16:1OH in *B. terrestris* and *B. lucorum* and of *Z*15-20:1OH in *B. lapidarius* can be ascribed to the absence of a FAR with the corresponding substrate specificity, but the very low amount of *Z*9-16:1 acyl in *B. lucorum* and *B. terrestris* male LG and the absence of detectable *Z*15-20:1 acyl in *B. lapidarius* LG also likely contribute (Supplementary Table 5).

We detected several fatty alcohols in FBs of *B. terrestris* and *B. lucorum*, 16:OH, *Z*9,*Z*12-18:2OH and *Z*9,*Z*12,*Z*15-18:3OH being the most abundant (Supplementary Table 6). Fatty alcohols are not expected to be transported from FB across haemolymph to LG^29^. However, the presence of *Z*9,*Z*12-18:2 and *Z*9,*Z*12,*Z*15-18:3 fatty alcohols in FB provides an explanation for why we did not find a FAR reducing *Z*9,*Z*12-18:2 and *Z*9,*Z*12,*Z*15-18:3 among the functionally characterized candidates from *B. terrestris* and *B. lucorum*. Our candidate selection criteria were based on the LG-specific FAR transcript abundance, and we might have disregarded a FAR that is capable of polyunsaturated fatty acyl reduction and is expressed at comparable levels in both LG and FB.

We noted several discrepancies between the FAR specificity in the yeast expression system and the apparent FAR specificity *in vivo* (i.e., the apparent specificity of fatty acyl reduction in bumblebee LG calculated from the fatty acyl and fatty alcohol content). We found that *Blap*FAR-A1 is capable of producing substantial amounts of *Z*9-16:1OH and *Z*9-18:1OH (Supplementary Fig. 7b, Supplementary Fig. 8) in the yeast system, while in *B. lapidarius* LG, only *Z*9-16:1 acyl is converted to *Z*9-16:1OH, as evidenced by the absence of detectable amounts of *Z*9-18:1OH (Fig. 6 and Supplementary Table 5). Additionally, *Blap*FAR-A1 and *Blap*FAR-A4 in the yeast expression system produce polyunsaturated fatty alcohols that are not present in *B. lapidarius* male LG, despite the presence of corresponding fatty acyls in the LG (Supplementary Table 5). A possible explanation for the differences between FAR specificities in the bumblebee LG and the yeast expression system is that the pool of LG fatty acyls that we assessed and used to evaluate the apparent FAR specificities has a different composition than the LG pool of fatty acyl-CoAs, which are the form of fatty acyls accepted by FARs as substrates. The relatively low concentrations of *Z*9,*Z*12-18:2CoA, *Z*9,*Z*12,*Z*15-18:3CoA, and Z9-18:1CoA in the LG of male *B. lapidarius* compared to the concentrations of the respective fatty acyls could prevent detectable accumulation of the corresponding fatty alcohols. We therefore propose that the selectivity of enzymes and binding proteins that convert fatty acyls to fatty acyl-CoAs^66^ and protect fatty acyl-CoAs from hydrolysis^67^ represents an additional mechanism shaping the species-specific fatty alcohol composition in bumblebee male LGs.

In sum, the functional characterization of bumblebee FARs indicates that the combined action of FARs from the expanded FAR-A orthology group has the capability to biosynthesize the majority of bumblebee MMP fatty alcohols. The substrate specificity of FARs apparently contributes to the species-specific MMP composition, but other biosynthetic steps, namely the process of fatty acyl and fatty acyl-CoA accumulation, likely also contribute to the final fatty alcohol composition of bumblebee MMPs.

## Conclusion

In the present work, we substantially broadened our limited knowledge of the function of FARs in Hymenoptera, one of the largest insect orders. The experimentally determined reductase specificity of FARs that are abundantly expressed in bumblebee male LGs is consistent with their role in MMP biosynthesis. The majority of these MMP-biosynthetic FARs belong to the FAR-A orthology group. We found that the FAR-A group expanded in the *Bombus* and Meliponini lineage. By conducting transcriptome- and genome-scale comparative studies of a FAR gene family across Hymenoptera, assaying tissue-specific FAR gene expression, and experimentally characterizing FAR enzymatic specificities, we provide evidence that lineage-specific gene family expansion shaped the genetic basis of pheromone production in the crown group of bees. Our analysis of TE distribution in the *B. terrestris* genome indicates that TEs enriched in the vicinity of FAR-A genes might have substantially contributed to the dramatic expansion of the FAR-A gene group. In the future, the increasing availability of genomic and transcriptomic resources for Hymenoptera should enable us to more precisely delineate the taxonomic extent and evolutionary timing of the massive FAR gene family expansion and assess in detail the role of TEs in the process.

## Methods

### Insects

Specimens of *Bombus lucorum* and *Bombus lapidarius* were obtained from laboratory colonies established from naturally overwintering bumblebee queens. The *Bombus terrestris* specimens originated from laboratory colonies obtained from a bumblebee rearing facility in Troubsko, Czech Republic.

LG and FB samples used for transcriptome sequencing were prepared from 3-day-old bumblebee males by pooling tissues from multiple specimens from the same colony. The cephalic part of the LG and a section of the abdominal peripheral FB were dissected, transferred immediately to TRIzol (Invitrogen), then flash-frozen at −80 °C and stored at this temperature prior to RNA isolation.

### RNA isolation and cDNA library construction

For cloning of FARs and RT-qPCR analysis of tissue-specific gene expression, RNA was isolated from individual bumblebee tissues by guanidinium thiocyanate-phenol-chloroform extraction followed by RQ1 DNase (Promega) treatment and RNA purification using the RNeasy Mini Kit (Qiagen). The tissue sample for RNA isolation from virgin queen LGs consisted of pooled glands from two specimens. For RT-qPCR analysis of age-specific expression in *B. lapidarius*, RNA was isolated using the Direct-zol RNA MicroPrep Kit (Zymo Research). A nanodrop ND-1000 spectrophotometer (Thermo Fisher) was employed to determine the isolated RNA concentration. The obtained RNA was kept at −80 °C until further use.

The cDNA libraries of LGs from 3-day-old bumblebee males were constructed from 0.50 μg total RNA using the SMART cDNA Library Construction Kit (Clontech) with either Superscript III (Invitrogen) or M-MuLV (New England Biolabs) reverse transcriptase.

### Transcriptome sequencing, assembly and annotation

The transcriptomes of male LGs and FBs of *B. lucorum* and *B. terrestris* were assembled as previously described^42^. The male LG and FB transcriptomes of *B. lapidarius* were sequenced and assembled as described^42^. Briefly, total RNA was isolated from the LGs and FBs of three 3-day-old *B. lapidarius* males and pooled into a FB and LG sample. Total RNA (5 µg) from each of the samples was used as starting material. Random primed cDNA libraries were prepared using poly(A)^+^ enriched mRNA and standard Illumina TrueSeq protocols (Illumina). The resulting cDNA was fragmented to an average of 150 bp. RNA-Seq was carried out by Fasteris (Fasteris) and was performed using an Illumina HiSeq 2500 Sequencing System. Quality control, including filtering high-quality reads based on the fastq score and trimming the read lengths, was carried out using CLC Genomics Workbench software v. 7.0.1 (http://www.clcbio.com). The complete transcriptome libraries were assembled *de novo* using CLC Genomics Workbench software. FAR expression values were calculated by mapping Illumina reads against the predicted coding regions of FAR sequences using bowtie2 v2.2.6^68^ and counting the mapped raw reads using htseq v0.9.1^69^. The raw read counts were normalized for the FAR coding region length and the total number of reads in the sequenced library, yielding reads per kilobase of transcript per million mapped reads (RPKM) values^70^. A constant value of 1 was added to each RPKM value and subsequently log2-transformed and visualized as heatmaps using the ggplot2 package in R. Complete short read (Illumina HiSeq2500) data for FB and LG libraries from *B. lucorum* and *B. lapidarius* were deposited in the Sequence Read Archive (https://www.ncbi.nlm.nih.gov/sra) with BioSample accession numbers SAMN08625119, SAMN08625120, SAMN08625121, and SAMN08625122 under BioProject ID PRJNA436452.

### FAR sequence prediction

The FARs of *B. lucorum* and *B. lapidarius* were predicted based on Blast2GO transcriptome annotation and their high protein sequence similarity to previously characterized FARs from the European honeybee *Apis mellifera*^26^ and the silk moth *Bombyx mori*^44^.

FARs from annotated genomes or transcriptomes of other hymenopteran species (*Bombus impatiens*^62^, *Bombus terrestris*^62^, *Melipona quadrifasciata*^71^, *Apis mellifera*^64^, *Megachile rotundata*^71^, *Dufourea novaeangliae, Camponotus floridanus*^72^, *Acromyrmex echinatior*^73^, *Harpegnathos saltator*^72^, *Nasoni vitripenis*^74^ and *Polistes canadensis*^75^) were retrieved by blastp searches (*E*-value cutoff 10^−5^) of the species-specific NCBI RefSeq protein database or UniProt protein database using predicted protein sequences of *B. lucorum, B. lapidarius* and *B. terrestris* FARs (accessed February 2017). An additional round of blastp searches using FARs found in the first blastp search round did not yield any additional significant (*E*-value < 10^−5^) blastp hits, indicating that all FAR homologs were found in the first round of blastp searches (data not shown). For FARs with multiple predicted splice variants, only the longest protein was used for phylogenetic tree reconstruction. FARs from non-annotated transcriptomes of *Bombus rupestris* and *Tetragonula carbonaria* were retrieved via tblastn search (*E*-value cutoff 10^−5^) of the publicly available contig sequences (BioProject PRJNA252240 and PRJNA252285, respectively^76^) using *Bombus* FARs as a query. The longest translated ORFs were used as a query in tblastn searches against NCBI non-redundant nucleotide database (nr/nt) and ORFs not yielding highly-scoring blast hits annotated as FARs were rejected. The final FAR proteins were used for gene tree reconstruction.

The active site, conserved Rossmann fold NAD(P)^+^ binding domain (NABD)^77^ and a putative substrate binding site in FAR coding sequences were predicted using Batch conserved domain search^78^. The matrix of protein identities was calculated using Clustal Omega with default parameters (https://www.ebi.ac.uk/Tools/msa/clustalo/accessedFebruary2018).

### FAR gene tree reconstruction

The protein sequences of predicted hymenopteran FARs were aligned using mafft v7.305. The gene tree was inferred in IQTREE v1.5.5 with 1,000 ultrafast bootstrap approximation replicates^79^, and with a model of amino acid substitution determined by ModelFinder^80^ implemented in IQTREE. The tree was visualized and annotated using the ggtree package^81^ in R programming language^82^.

### Genome alignment and TE-enrichment analysis

The genomes of *A. mellifera* and *B. terrestris* were aligned using MAUVE 2.4.0^83^. The genomic position of predicted *B. terrestris* FAR genes was visually inspected using the NCBI Graphical sequence viewer (accessed January 2018 at Nucleotide Entrez Database).

TE-enrichment analysis in the vicinity of FAR genes in the *B. terrestris* genome was carried out to explore the impact of TEs in extensive expansion of FAR-A genes. The genomic data were retrieved from the FTP server of the Bumble Bee Genome Project (accessed March 2018)^62^. TE density around FAR genes was calculated 10 kb upstream and downstream of each FAR gene, separately for FAR-A genes and non-FAR-A genes. Statistical significances were obtained by permutation test. We compared FAR-A/non-FAR-A gene set average TE density to the null distribution of the average TE densities around *B. terrestris* genes built by 10,000 randomly sampling gene sets of a size corresponding to the size of the FAR-A/non-FAR-A gene set from publicly available Augustus gene predictions^62^. TE densities were analysed for pooled set of all TEs and separately for each TE class and major TE family (Class I: LINE, LTR, LARD; Class II: TIR, MITE, TRIM) using custom shell scripts and bedtools, a suite of Unix genomic tools^84^. R programming language was used for statistical analysis^82^.

### Quantitative PCR analysis of FAR expression

First-strand cDNA was synthesized from 0.30 μg total RNA using oligo(dT)_12-18_ primers and Superscript III reverse transcriptase. The resulting cDNA samples were diluted 5-fold with water prior to RT-qPCR. The primers used for the assay (Supplementary Table 4) were designed with Primer-BLAST (https://www.ncbi.nlm.nih.gov/tools/primer-blast/)^85^ and first tested for specificity by employing amplicon melting curve analysis on pooled cDNAs from each species.

The reaction mixtures were prepared in a total volume of 20 μL consisting of 2 μL sample and 500 nM of each primer using LightCycler 480 SYBR Green I Master kit (Roche). The reactions were run in technical duplicates for each sample. RT-qPCR was performed on a LightCycler 480 Instrument II (Roche) in 96-well plates under the following conditions: 95 °C for 60 s, then 45 cycles of 95 °C for 30 s, 55 °C for 30 s and 72 °C for 30 s followed by a final step at 72 °C for 2 min.

The acquired data were processed with LightCycler 480 Software 1.5 (Roche) and further analysed with MS Excel (Microsoft Corporation). FAR transcript abundances were normalized to the reference genes phospholipase A2 (PLA2) and elongation factor 1α (eEF1α) as described^86^.

### FAR gene isolation and cloning

The predicted coding regions of FARs from *B. lucorum, B. lapidarius* and *B. terrestris* were amplified by PCR from LG cDNA libraries using gene-specific primers (Supplementary Table 4) and Phusion HF DNA polymerase (New England Biolabs). Parts of the full-length coding sequence of *Blap*FAR-A5 were obtained by RACE procedure. Briefly, the PCR-amplified sequences containing the 5’ and 3’ ends of *Blap*FAR-A5 were inserted into pCR2.1 TOPO vector using TOPO TA Cloning kit (Invitrogen) and sequenced by Sanger method. The resulting sequences overlapped with contig sequences retrieved from the *B. lapidarius* transcriptome. The full-length *Blap*FAR-A5-coding region was subsequently isolated using gene-specific PCR primers. The sequence of *Blap*FAR-A1 and yeast codon-optimized sequences of *Bluc*FAR-A1-opt and *Bluc*FAR-A2-opt were obtained by custom gene synthesis (Genscript); see Supplementary Data 1 for synthetic sequences. The individual FAR coding regions were then inserted into linearized pYEXTHS-BN vector^87^ using the following restriction sites: *Bter*/*Bluc*FAR-A1 and *Blap*FAR-J at *Sph*I-*Not*I sites; *Bter*/*Bluc*FAR2, *Blap*FAR-A1, *Blap*FAR-A1-short and *Blap*FAR-A5 at *Bam*HI-*Not*I sites; and *Bluc*FAR-A1-opt/FAR-A2-opt and *Blap*FAR-A4 at *Bam*HI-*Eco*RI sites. In the case of *Bter*FAR-J, the *Taq* DNA polymerase (New England Biolabs)-amplified sequence was first inserted into pCR2.1 TOPO vector and then subcloned into pYEXTHS-BN via *Bam*HI-*Eco*RI sites using the In-Fusion HD Cloning kit (Clontech).

The resulting vectors containing FAR sequences *N*-terminally fused with 6×His-tag were subsequently transformed into *E. coli* DH5α cells (Invitrogen). The plasmids were isolated from bacteria with Zyppy Plasmid Miniprep kit (Zymo Research) and Sanger sequenced prior to transformation into yeast. The protein-coding sequences of all studied FARs were deposited to GenBank under accession numbers MG450697‒MG450704 and MG930980‒MG930983.

### Functional assay of FARs in yeast

Expression vectors carrying FAR-coding sequences were transformed into *Saccharomyces cerevisiae* strain BY4741 (*MAT*a *his3*Δ*1 leu2*Δ*0 met15*Δ*0 ura3*Δ*0*)^88^ using S.c. EasyComp Transformation Kit (Invitrogen). To test FAR specificity, yeasts were cultured for 3 days in 20 mL synthetic complete medium lacking uracil (SC−U) supplemented with 0.5 mM Cu^2+^ (inducer of heterologous gene expression), 0.2% peptone and 0.1% yeast extract. The yeast cultures were then washed with water and the cell pellets lyophilized before proceeding with lipid extraction. FAR specificities were determined with the FARs acting on natural substrates present in yeast cells and with individual fatty acyls added to the cultivation media, with the respective fatty alcohols present in the LGs of studied bumblebees. Yeast cultures were supplemented with the following fatty acyls: 0.1 mM *Z*9,*Z*12-18:2COOH (linoleic acid), *Z*9,*Z*12,*Z*15-18:3COOH (*α*-linolenic acid) or *Z*15-20:1Me solubilized with 0.05% tergitol. We chose *Z*15:20:1 as a representative monounsaturated C20+ fatty acyl substrate because *Z*15-20:1OH is the most abundant monounsaturated fatty alcohol in *B. terrestris* LG (Fig. 6 and Supplementary Table 5).

The level of heterologous expression of bumblebee FARs was assayed by Western blot analysis of the whole-cell extracts (obtained via sonication) using anti-6×His-tag antibody-HRP conjugate (Sigma-Aldrich) and SuperSignal West Femto Maximum Sensitivity Substrate kit (Thermo Fisher Scientific).

### Lipid extraction and transesterification

Lipids were extracted from bumblebee tissue samples under vigorous shaking using a 1:1 mixture of CH_2_Cl_2_/MeOH, followed by addition of an equal amount of hexane and sonication. The extracts were kept at −20 °C prior to GC analysis.

Base-catalysed transesterification was performed as described previously^89^ with modifications: the sample was shaken vigorously with 1.2 mL 2:1 CH_2_Cl_2_/MeOH and glass beads (0.5 mm) for 1 h. After brief centrifugation to remove particulate debris, 1 mL supernatant was evaporated under nitrogen, and the residue was dissolved using 0.2 mL 0.5 M KOH in methanol. The mixture was shaken for 0.5 h and then neutralized by adding 0.2 mL Na_2_HPO_4_ and KH_2_PO_4_ (0.25 M each) and 35 μL 4 M HCl. The obtained FAMEs were extracted with 600 μL hexanes and analysed by gas chromatography.

For quantification purposes, either 1-bromodecane (10:Br) or 1-bromoeicosane (20:Br) were added to the extracts as internal standards.

### Gas chromatography and fatty alcohol ratio determination

Standards of *Z*9,*Z*12,*Z*15-18:3OH and *Z*15-20:1OH were prepared from their corresponding acids/FAMEs by reduction with LiAlH_4_. The *Z*9-18:1Me standard was prepared by reacting oleoyl chloride with methanol. Other FAME and fatty alcohol standards were obtained from commercial suppliers. The FA-derived compounds in extracts were identified based on the comparison of their retention times with the standards and comparison of measured MS spectra with those from spectral libraries. Double bond positions were assigned after derivatization with dimethyl disulfide^90^.

The fatty alcohol ratio was calculated as the molar percentage of fatty alcohol relative to the total fatty alcohol and fatty acyls (e.g. free FAs, fatty acyl-CoAs, and triacylglycerols) containing the same fatty chain structure, i.e., the same chain length and double bond position. The fatty alcohol ratio represents the hypothetical degree of conversion of total fatty acyls (all the quantified fatty acyls counted as fatty acyl-CoAs) to the respective fatty alcohol and reflects the apparent FAR specificity in the investigated bumblebee tissue or yeast cell.

### GC-FID

A flame-ionization detector was used for quantitative assessment of the FA-derived compounds. The separations were performed on a Zebron ZB-5ms column (30 m × 250 μm I. D. × 0.25 μm film thickness, Phenomenex) using a 6890 gas chromatograph (Agilent Technologies) with following parameters: helium carrier gas, 250 °C injector temperature, and 1 mL.min^−1^column flow. The following oven temperature program was used: 100 °C (held for 1 min), ramp to 285 °C at a rate of 4 °C.min^−1^ and a second ramp to 320 °C at a rate of 20 °C.min^−1^ with a final hold for 5 min at 320 °C. The analytes were detected in FID at 300 °C using a makeup flow of 25 mL.min^−1^ (nitrogen), hydrogen flow of 40 mL.min^−1^, air flow of 400 mL.min^−1^ and acquisition rate of 5 Hz. The collected data were processed in Clarity (DataApex).

### GC×GC-MS

This approach was used to identify analytes by comparing their retention characteristics and mass spectra with those of synthetic standards. The following conditions were employed using a 6890N gas chromatograph (Agilent Technologies) coupled to a Pegasus IV D time-of-flight (TOF) mass selective detector (LECO Corp.): helium carrier gas, 250 °C injector temperature, 1 mL.min^−1^column flow, modulation time of 4 s (hot pulse time 0.8 s, cool time 1.2 s), modulator temperature offset of +20 °C (relative to secondary oven) and secondary oven temperature offset of +10 °C (relative to primary oven). Zebron ZB-5ms (30 m × 250 μm I. D. × 0.25 μm film thickness, Phenomenex) was used as a non-polar primary column and BPX-50 (1.5 m × 100 μm I. D. × 0.10 μm film thickness, SGE) was used as a more polar secondary column. The primary oven temperature program was as follows: 100 °C (1 min), then a single ramp to 320 °C at a rate of 4 °C.min^−1^ with a final hold for 5 min at 320 °C.

The mass selective detector was operated in electron ionization mode (electron voltage −70 V) with a transfer line temperature of 260 °C, ion source temperature of 220 °C, 100 Hz acquisition rate, mass scan range of 30–600 u and 1800 V detector voltage. ChromaTOF software (LECO Corp.) was used to collect and analyse the data.

### Organic synthesis

Methyl *Z*15-eicosenoate (*Z*15-20:1Me, **4**) was synthesized by a new and efficient four-step procedure, starting from inexpensive and easily available cyclopentadecanone. The C1–C15 part of the molecule was obtained by Baeyer-Villiger oxidation of cyclopentadecanone, followed by subsequent methanolysis of the resulting lactone **1** and Swern oxidation of the terminal alcohol group of **2**; the C16–C20 fragment was then connected to the aldehyde **3** by Wittig olefination.

All reactions were conducted in flame- or oven-dried glassware under an atmosphere of dry nitrogen. THF, CH_2_Cl_2_ and MeOH were dried following standard methods under a nitrogen or argon atmosphere. Petroleum ether (PE, 40–65 °C boiling range) was used for chromatographic separations. TLC plates (silica gel with fluorescent indicator 254 nm, Fluka or Macherey-Nagel) were used for reaction monitoring. Flash column chromatographic separations were performed on silica gel 60 (230–400 mesh, Merck or Acros).

IR spectra were taken on an ALPHA spectrometer (Bruker) as neat samples using an ATR device. ^1^H and ^13^C NMR spectra were recorded in CDCl_3_ on an AV III 400 HD spectrometer (Bruker) equipped with a cryo-probe or an AV III 400 spectrometer (Bruker) equipped with an inverse broad-band probe at 400 MHz for ^1^H and 100 MHz for ^13^C. ^1^H NMR chemical shifts were provided in ppm using TMS as external standard; ^13^C NMR chemical shifts were referenced against the residual solvent peak. The connectivity was determined by ^1^H-^1^H COSY experiments. GC-MS (EI) measurements were performed on an Agilent 5975B MSD coupled to a 6890N gas chromatograph (Agilent Technologies). High-resolution MS (HRMS) spectra were measured on a Q-Tof micro spectrometer (resolution 100000 (ESI), Waters) or GCT Premier orthogonal acceleration TOF mass spectrometer (EI and CI, Waters).

### 1-Oxacyclohexadecan-2-one (1)

Cyclopentadecanone (500 mg, 2.23 mmol) was dissolved in dry CH_2_Cl_2_ (6 mL) and *meta*-chloroperbenzoic acid (*m*CPBA) (687 mg, 2.79 mmol, 70%) was added at 0 °C. The reaction mixture was stirred at room temperature (r.t.), occasionally concentrated under a flow of nitrogen, and the solid residue was re-dissolved in dry CH_2_Cl_2_. After stirring for four days, the conversion was still not complete; additional *m*CPBA (164 mg, 667 μmol, 70%) was added at 0 °C and stirring was continued at r.t. for 48 h. The mixture was diluted with CH_2_Cl_2_ (20 mL), and the organic layer was washed with saturated NaHCO_3_ solution (5×5 mL) and brine (5 mL). The organic layer was dried over Na_2_SO_4_, filtered and evaporated. The crude product was purified by column chromatography (50 mL silica gel, PE/CH_2_Cl_2_ 1:1) providing product **1** (426 mg, 80%) as a colorless waxy solid.

**1**: Melting point (m.p.) <30 °C. *R*_f_ (PE/Et_2_O 95:5) = 0.5. IR (film): *v* = 2925, 2855, 1733, 1459, 1385, 1349, 1234, 1165, 1108, 1070, 1013, 963, 801, 720 cm^−1^. HRMS (+EI TOF) *m*/*z*: (C_15_H_28_O_2_) calc.: 240.2089, found: 240.2090. ^1^H NMR (400 MHz, CDCl_3_) *δ* = 4.13 (t, *J* = 5.7 Hz, 2H, H16), 2.33 (t, *J* = 7.0 Hz, 2H, H3), 1.72–1.56 (m, 4H, H4, H15), 1.48–1.37 (m, 2H, H14), 1.36–1.23 (m, 18H, H5–H13). ^13^C NMR (100 MHz, CDCl_3_) *δ* = 174.2, 64.1, 34.6, 28.5, 27.9, 27.28, 27.26, 27.1, 26.8, 26.5, 26.2, 26.1, 26.0, 25.3, 25.1.

### Methyl 15-hydroxypentadecanoate (2)

MeOK (74 μL, 208 μmol, 2.81M in MeOH) was added dropwise at 0° C to a mixture of lactone **1** (50 mg, 208 μmol), dry THF (0.5 mL) and MeOH (1 mL). The mixture was stirred at r.t. for 48 h, by which point the reaction was complete as indicated by TLC. The solution was quenched with a few drops of water and diluted with Et_2_O (5 mL). After stirring for 30 min, the layers were separated and the aqueous layer was extracted with Et_2_O (3×3 mL). The combined organic layers were washed with brine and water, dried over Na_2_SO_4_, filtered and evaporated to obtain nearly pure product. Purification by column chromatography (5 mL silica gel, PE/EtOAc 9:1) provided product **2** (55 mg, 97%) as a colorless solid.

**2**: m.p. 47–48 °C. *R*_f_ (PE/Et_2_O 95:5) = 0.2. IR (film): *v* = 3285, 2917, 2849, 1740, 1473, 1463, 1435, 1412, 1382, 1313, 1286, 1264, 1240, 1217, 1196, 1175, 1117, 1071, 1061, 1049, 1025, 1013, 992, 973, 926, 884, 731, 720, 701 cm^−1^. HRMS (+ESI) *m*/*z*: (C_16_H_32_O_3_Na) calc.: 295.2244, found: 295.2245. ^1^H NMR (400 MHz, CDCl_3_) *δ* = 3.64 (s, 3H, OCH_3_), 3.61 (t, *J* = 6.7 Hz, 2H, H15), 2.28 (t, *J* = 7.5 Hz, 2H, H2), 1.72 (s, 1H, OH), 1.59 (quint, *J* = 7.1 Hz, 2H, H3), 1.54 (quint, *J* = 7.1 Hz, 2H, H14), 1.37–1.14 (m, 20H, H4–H13). ^13^C NMR (101 MHz, CDCl_3_) *δ* = 174.5, 63.1, 51.6, 34.2, 32.9, 29.71 (3C), 29.68 (2C), 29.5 (2C), 29.4, 29.3, 25.9, 25.1.

### Methyl 15-oxopentadecanoate (3)

Dry DMSO (110 μL, 1.54 mmol) was added at −78 °C dropwise to a mixture of oxalyl chloride (90 μL, 1.03 mmol) and CH_2_Cl_2_ (2 mL) in a 25 mL flask, and the reaction mixture was stirred for 15 min. The hydroxy ester **2** (140 mg, 0.51 mmol) in dry CH_2_Cl_2_ (2 mL) was added dropwise via a cannula; the white, turbid reaction mixture was stirred for 40 min, and dry triethylamine (432 μL, 3.08 mmol) was added dropwise. The mixture was stirred at –78 °C for 1 h and warmed to 0 °C over 30 min at which point the reaction was complete according to TLC. The reaction mixture was diluted with CH_2_Cl_2_ (10 mL), quenched with saturated NH_4_Cl solution (5 mL) and water (5 mL), and warmed to r.t. The layers were separated and the aqueous layer was extracted with CH_2_Cl_2_ (3×10 mL). The combined organic layers were washed with brine, dried over MgSO_4_, filtered and evaporated. The crude product was purified by flash chromatography (10 mL silica gel, PE/EtOAc 95:5) giving aldehyde **3** (109 mg, 78%) as a colorless waxy solid.

**3**: m.p. <37 °C. *R*_f_ (PE/EtOAc 9:1) = 0.4. IR (film): *v* = 2923, 2852, 2752, 1738, 1465, 1436, 1362, 1315, 1243, 1197, 1172, 1120, 1017, 985, 958, 883, 811, 719 cm^−1^. HRMS (+CI TOF) *m*/*z*: (C_16_H_31_O_3_) calc.: 271.2273, found: 271.2277. 1H NMR (400 MHz, CDCl_3_) *δ* = 9.76 (t, *J* = 1.9 Hz, 1H, H15), 3.66 (s, 3H, OCH_3_), 2.41 (td, *J* = 7.4, 1.9 Hz, 2H, H14), 2.29 (t, *J* = 7.5 Hz, 2H, H2), 1.67–1.56 (m, 4H, H3,H13), 1.35–1.20 (m, 18H, H4-H12). ^13^C NMR (100 MHz, CDCl_3_) *δ* = 203.1, 174.5, 51.6, 44.1, 34.3, 29.72, 29.70 (2C), 29.58, 29.56, 29.5, 29.4, 29.31, 29.29, 25.1, 22.2.

### Methyl *Z*15-eicosenoate (4, *Z*15-20:1Me)

NaHMDS (614 μL, 0.614 mmol, 1.0 M in THF) was added dropwise at −55 °C over 10 min to a suspension of high vacuum-dried (pentyl)triphenylphosphonium bromide (282 mg, 0.68 mmol)^91^ in dry THF (3 mL) in a flame dried round-bottomed Schlenk flask. The bright orange reaction mixture was stirred while warming to −40 °C for 50 min, and a solution of aldehyde **3** (92 mg, 0.34 mmol) in dry THF (1.5 mL) was added dropwise via cannula at −45 °C. Stirring was continued for 1 h, and the reaction mixture was warmed to r.t. over 90 min. The reaction mixture was diluted with PE (25 mL); filtered through a short silica gel plug, which was washed with PE; and evaporated. The crude product was purified by flash chromatography (silica gel, gradient PE/EtOAc 100:0 to 95:5) to give methyl ester **4** (88 mg, 79%) as a colorless oil.

**4**: *R*_f_ (PE/Et_2_O 95:5) = 0.6. IR (film): *v* = 3005, 2922, 2853, 1743, 1699, 1684, 1653, 1541, 1521, 1507, 1489, 1436, 1362, 1196, 1169, 1106, 1017, 880, 722 cm^−1^. GC-MS (EI) *t*_R_ [60 °C (4 min) → 10 °C/min to 320 °C (10 min)] 21 min; *m*/*z* (%): 324 (4) [M^+^], 292 (26), 250 (10), 208 (9), 152 (7), 123 (12), 111 (22), 97 (48), 87 (40), 83 (52), 74 (56), 69 (70), 59 (14), 55 (100), 41 (43), 28 (26). HRMS (+EI TOF) *m*/*z*: (C_21_H_40_O_2_) calc.: 324.3028, found: 324.3026. ^1^H NMR (401 MHz, CDCl_3_) *δ* = 5.39–5.30 (m, 2H, H15,H16), 3.66 (s, 3H, OCH_3_), 2.30 (t, *J* = 7.5 Hz, 2H, H2), 2.07–1.96 (m, 4H, H14,H17), 1.61 (quint, *J* = 7.5 Hz, 2H, H3), 1.37–1.14 (m, 24H, H4-H13,H18,H19), 0.89 (t, *J* = 7.2 Hz, 3H, H20). ^13^C NMR (100 MHz, CDCl_3_) *δ* = 174.5, 130.1, 130.0, 51.6, 34.3, 32.1, 29.9, 29.81, 29.79, 29.74, 29.70, 29.68, 29.6, 29.5, 29.4, 29.3, 27.4, 27.1, 25.1, 22.5, 14.2.

### Statistical analysis

All lipid quantifications in yeast and LGs and FBs of bumblebees and transcript quantifications in bumblebee tissues were performed using three biological replicates (in addition, technical duplicates were used for RT-qPCR). The results are reported as mean value ± S.D. Significant differences were determined by one-way analysis of variance (ANOVA) followed by post-hoc Tukey’s honestly significant difference (HSD) test or by a two-tailed *t*-test as indicated in the Results section.

## Acknowledgements

This work was supported by the Czech Science Foundation (grant no. 15-06569S) and by the Ministry of Education of the Czech Republic (project n. LO1302). We thank Dr. Stefan Jarau for stimulating discussions. Computational resources were provided by the CESNET LM2015042 and the CERIT Scientific Cloud LM2015085, provided under the program “Projects of Large Research, Development, and Innovations Infrastructures” and by the Okinawa Institute of Science and Technology.

## Supplementary Materials

**Supplementary Figure 1.**
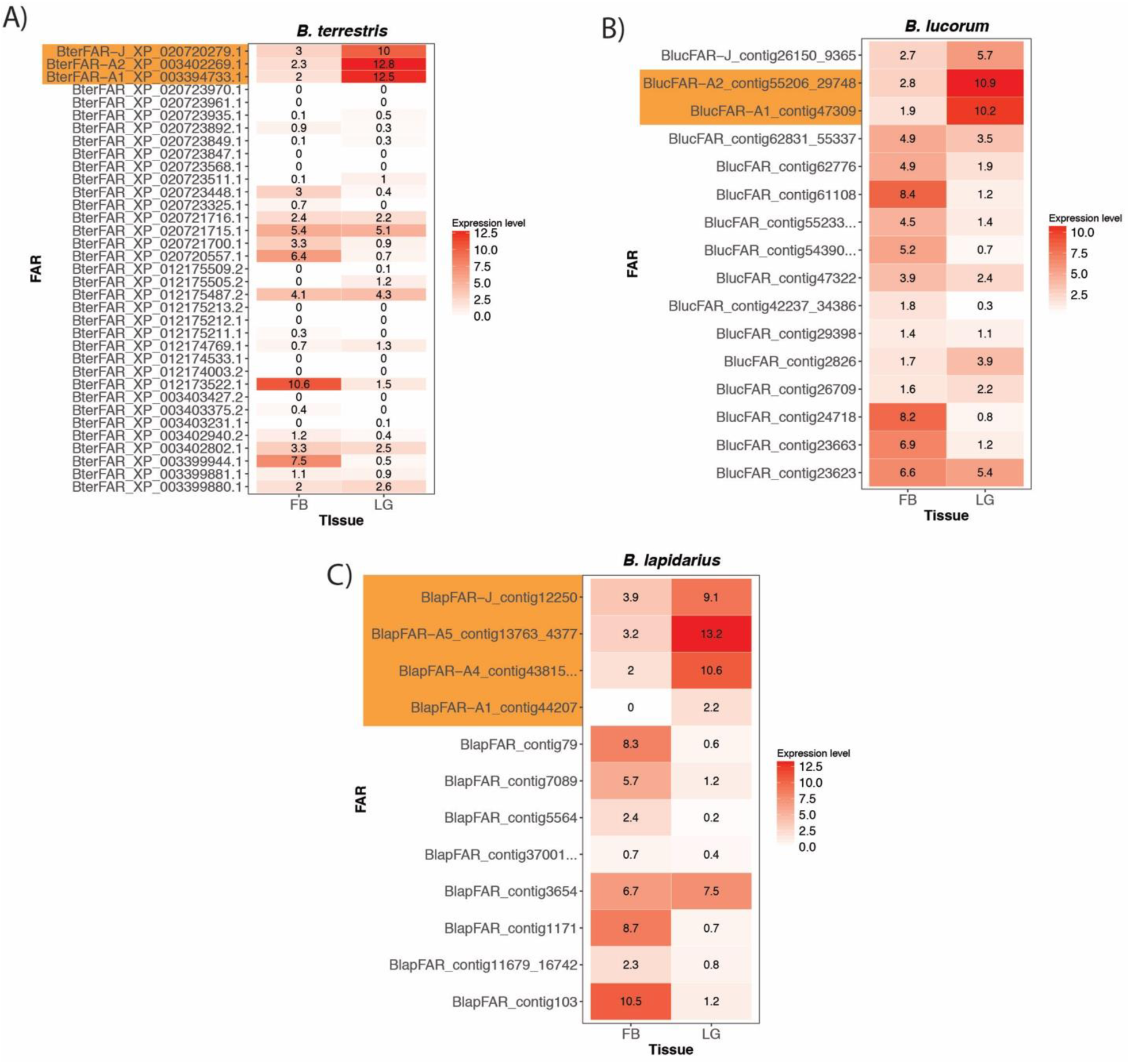
Expression of FARs in male labial gland and male fat body of *Bombus terrestris* (**A**), *B. lucorum* (**B**) and *B. lapidarius* (**C**). The expression values shown are log2-transformed normalized counts of reads (RPKM values) mapping to the FAR coding regions. FARs functionally characterized in this study are highlighted in orange.

**Supplementary Figure 2.**
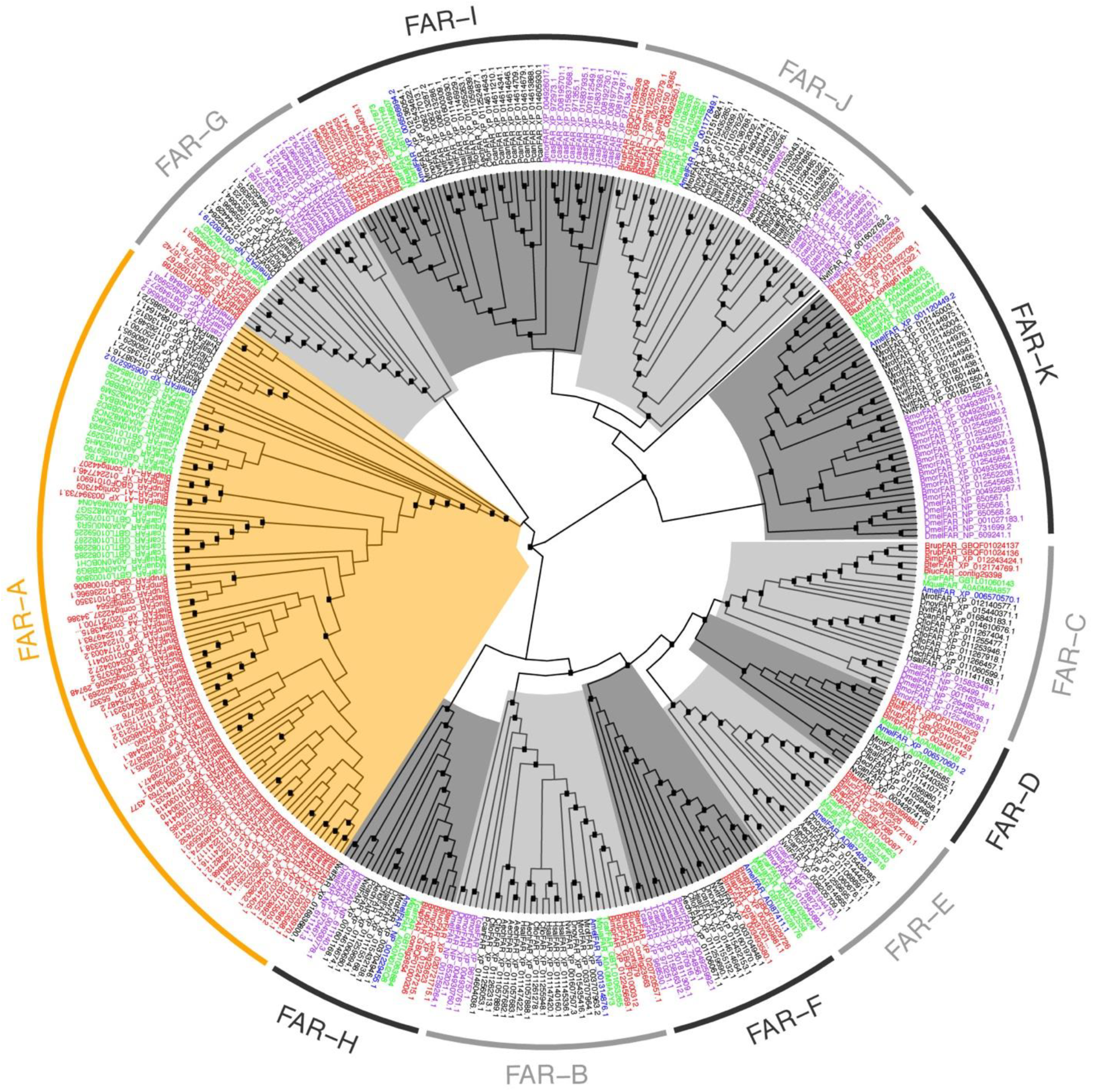
FAR gene tree including non-hymenopteran (*Drosophila melanogaster, Bombyx mori, Tribolium castaneum*) and hymenopteran FARs. Tree tips are coloured according to taxonomy: red, bumblebee FARs (*B. terrestris, B. lucorum, B. lapidarius, B. impatiens, B. rupestris*); green, stingless bee FARs (*Tetragonula carbonaria, Melipona quadrifasciata*); blue, *A. mellifera* FARs; black, FARs from other hymenopteran species; purple, non-hymenopteran species. The FAR-A orthology group is highlighted in orange; other orthologous groups are in shades of grey. Internal nodes highlighted with black boxes indicate bootstrap support >95 %.

**Supplementary Figure 3.**
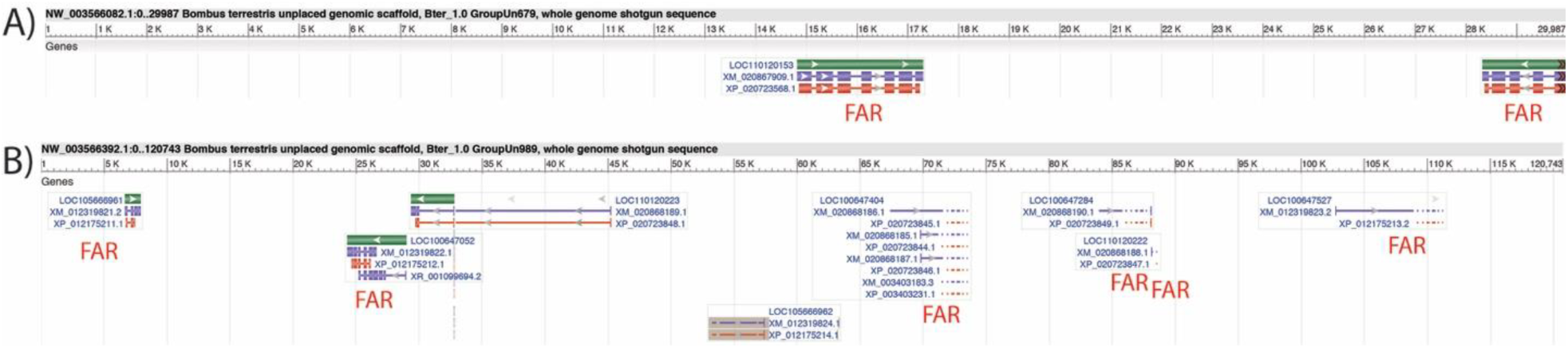
Clusters of *B. terrestris* FAR-A genes on *B. terrestris* genomic scaffolds Un679 (**A**) and Un989 (**B**). The horizontal axis shows genomic coordinates of genes within the scaffold. FAR genes are labelled.

**Supplementary Figure 4.**
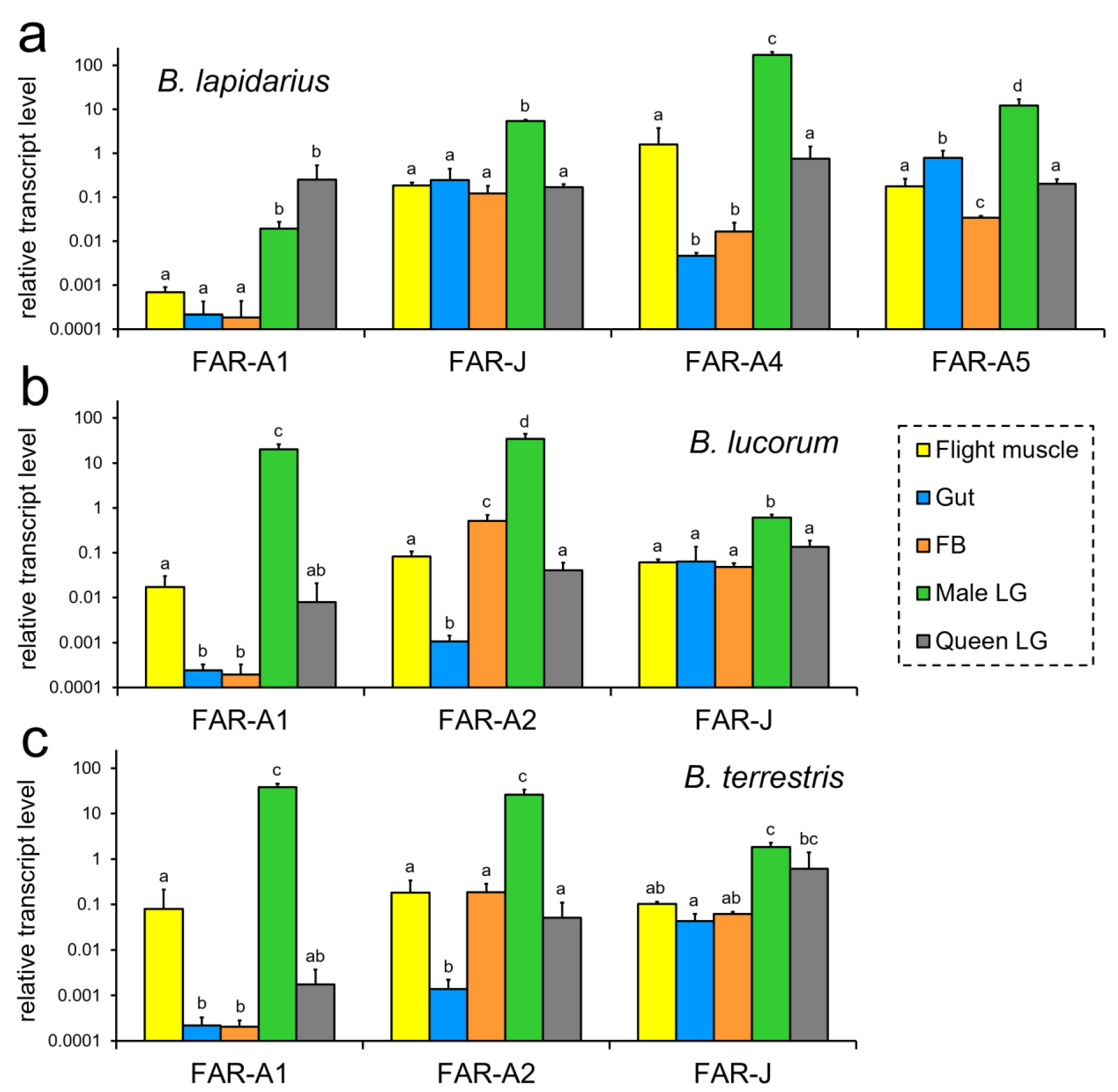
Relative transcript levels of FARs in the tissues of *B. lapidarius* (**a**), *B. lucorum* (**b**) and *B. terrestris* (**c**). Significant differences (*p* < 0.05, one-way ANOVA followed by *post*-*hoc* Tukey’s HSD test) are marked with different letters.

**Supplementary Figure 5.**
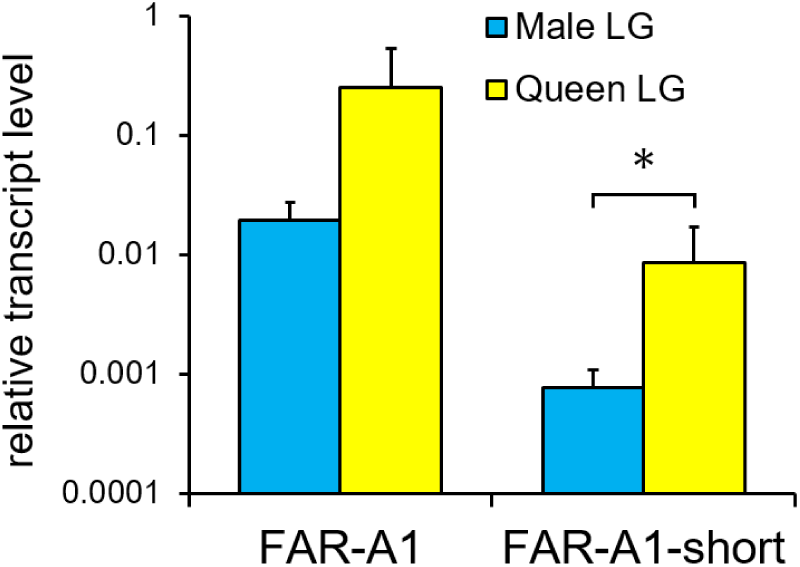
Relative transcript levels of FAR1 and FAR1-short in male and queen LGs of *B. lapidarius*. Significant differences (*p* < 0.05, two-tailed *t*-test) are indicated with an asterisk.

**Supplementary Figure 6.**
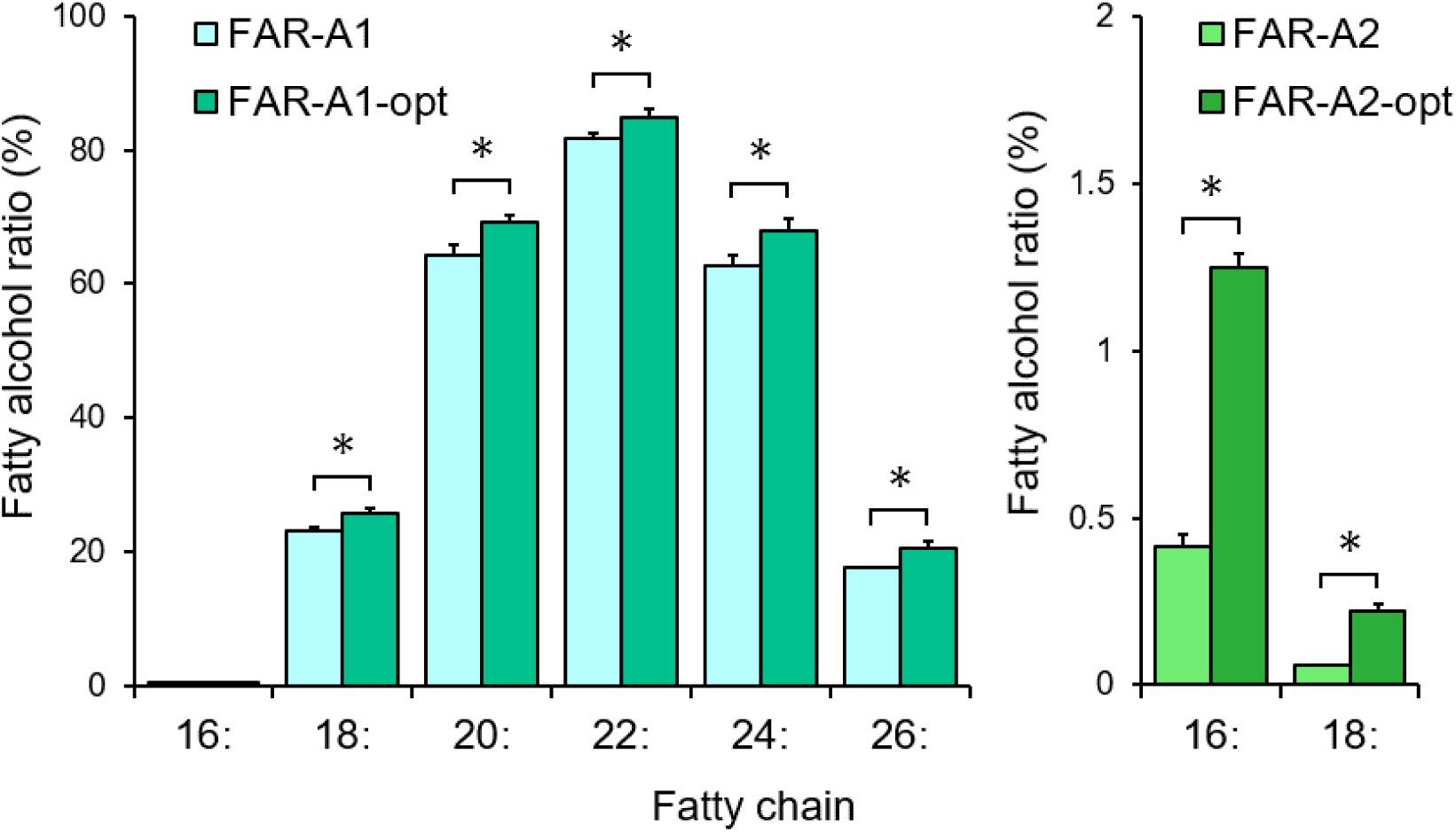
Fatty alcohol production by FARs from *B. lucorum* expressed from yeast codon-optimized and wild-type nucleotide sequences. Significant differences (*p* < 0.05, two-tailed *t*-test) are marked with asterisks. See Methods for a description of fatty alcohol ratio calculation.

**Supplementary Figure 7.**
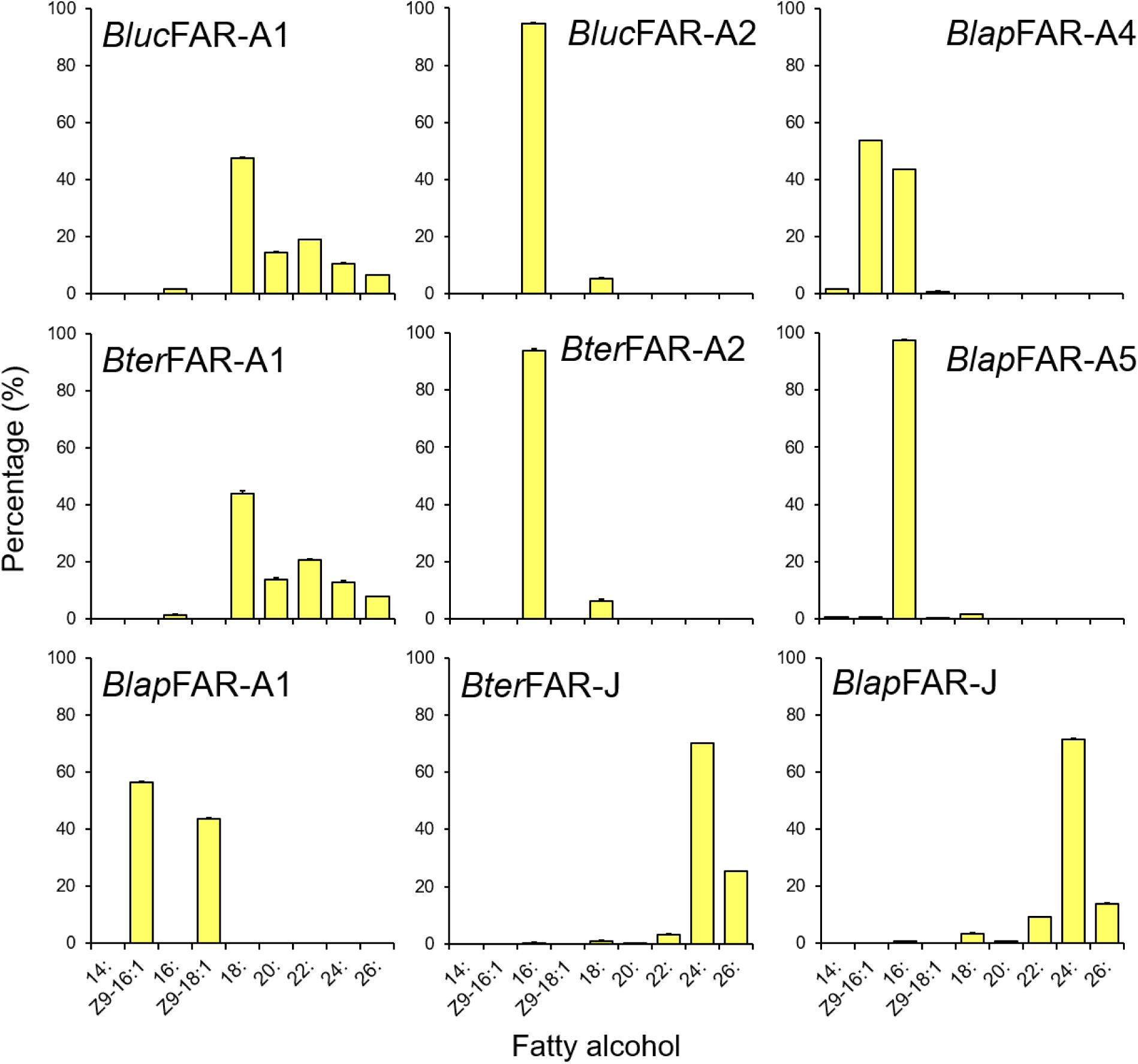
Fatty alcohol production in yeast strains expressing bumblebee FARs calculated as ratios of a particular alcohol to the total produced fatty alcohols.

**Supplementary Figure 8.**
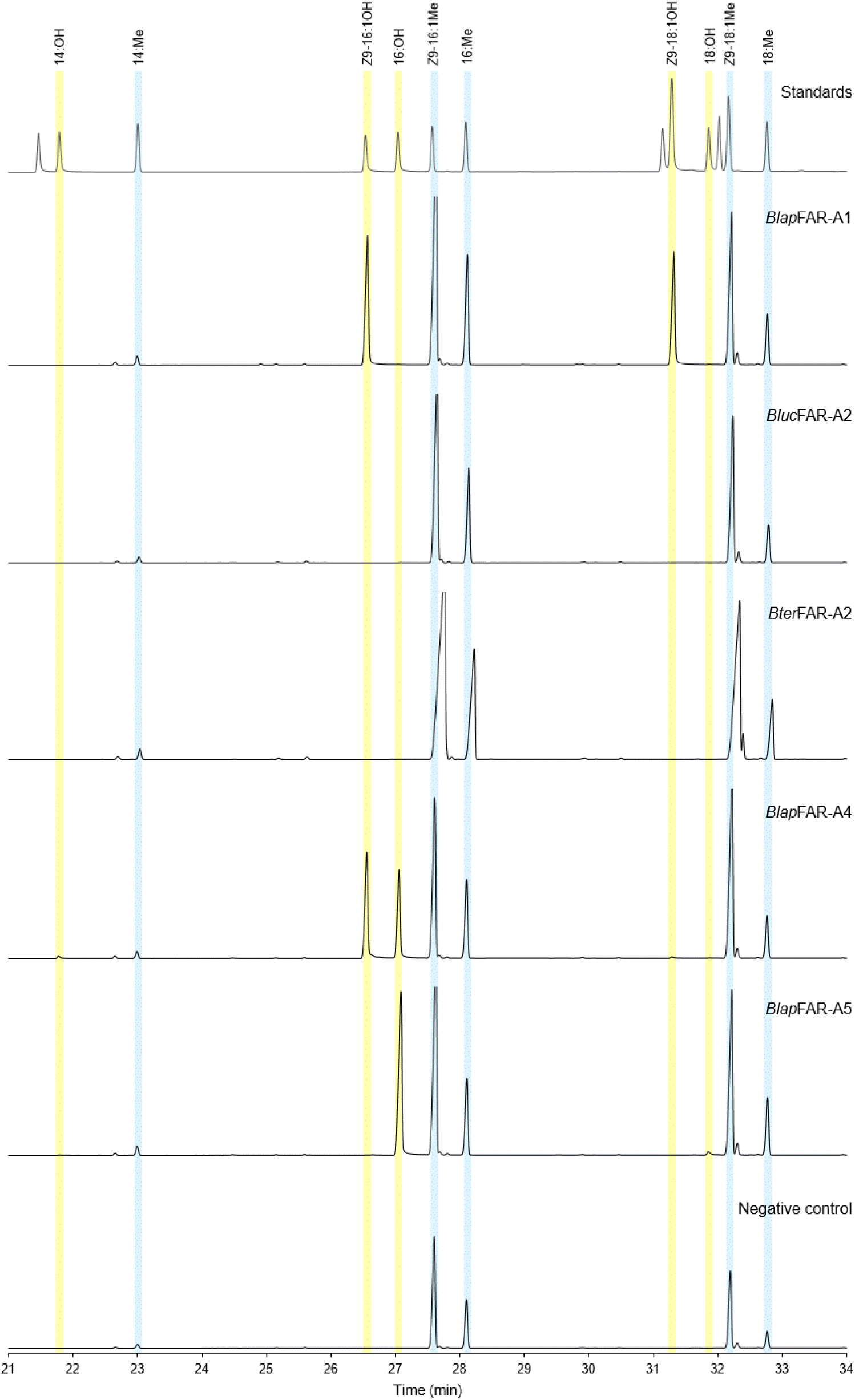
GC-FID chromatograms of extracts from control yeast strain and strains expressing FARs specific for long chain fatty acyls (C14–C18).

**Supplementary Figure 9.**
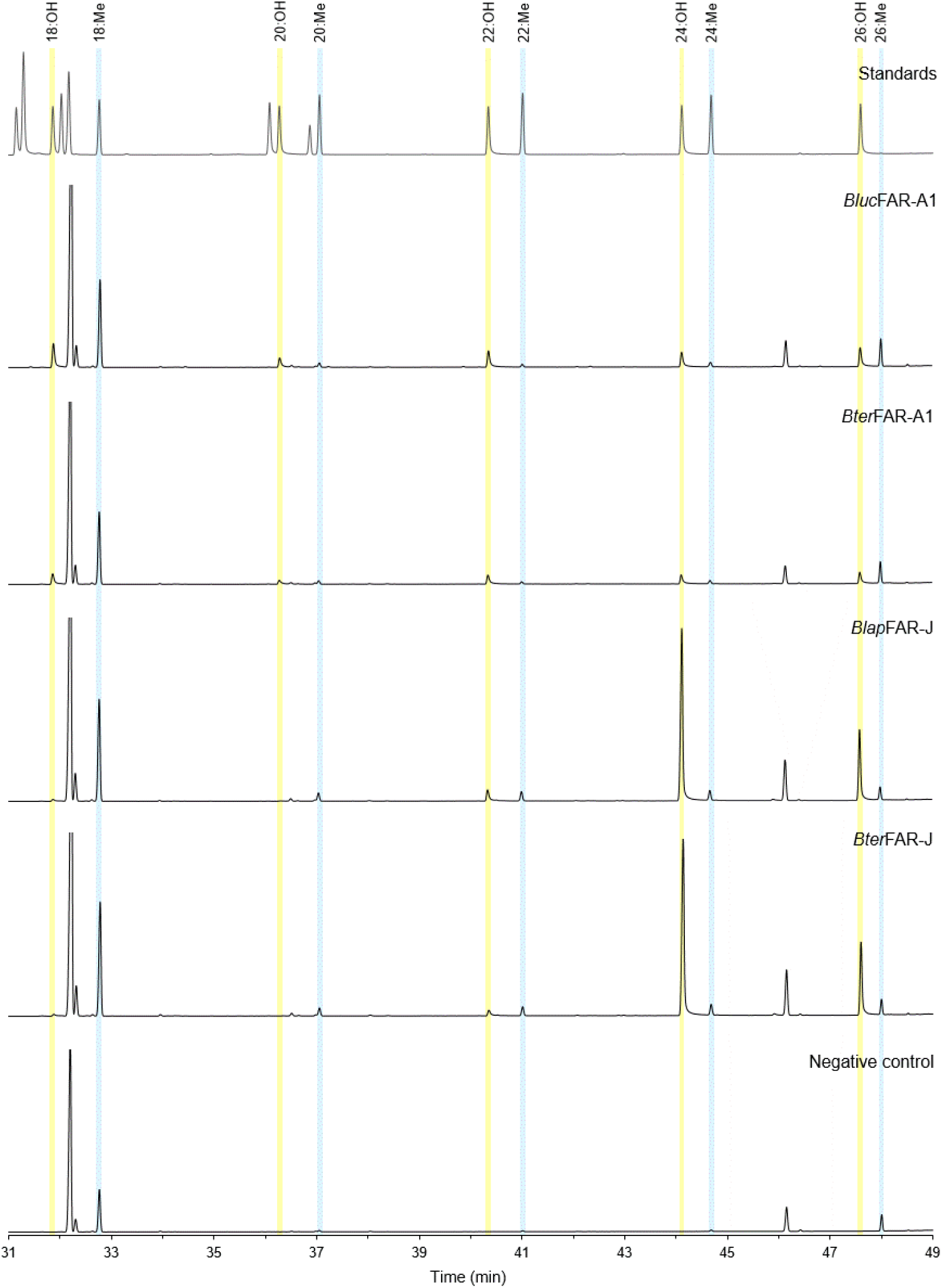
GC-FID chromatograms of extracts from control yeast strain and strains expressing FARs specific for very long chain fatty acyls (C20–C26).

**Supplementary Figure 10.**
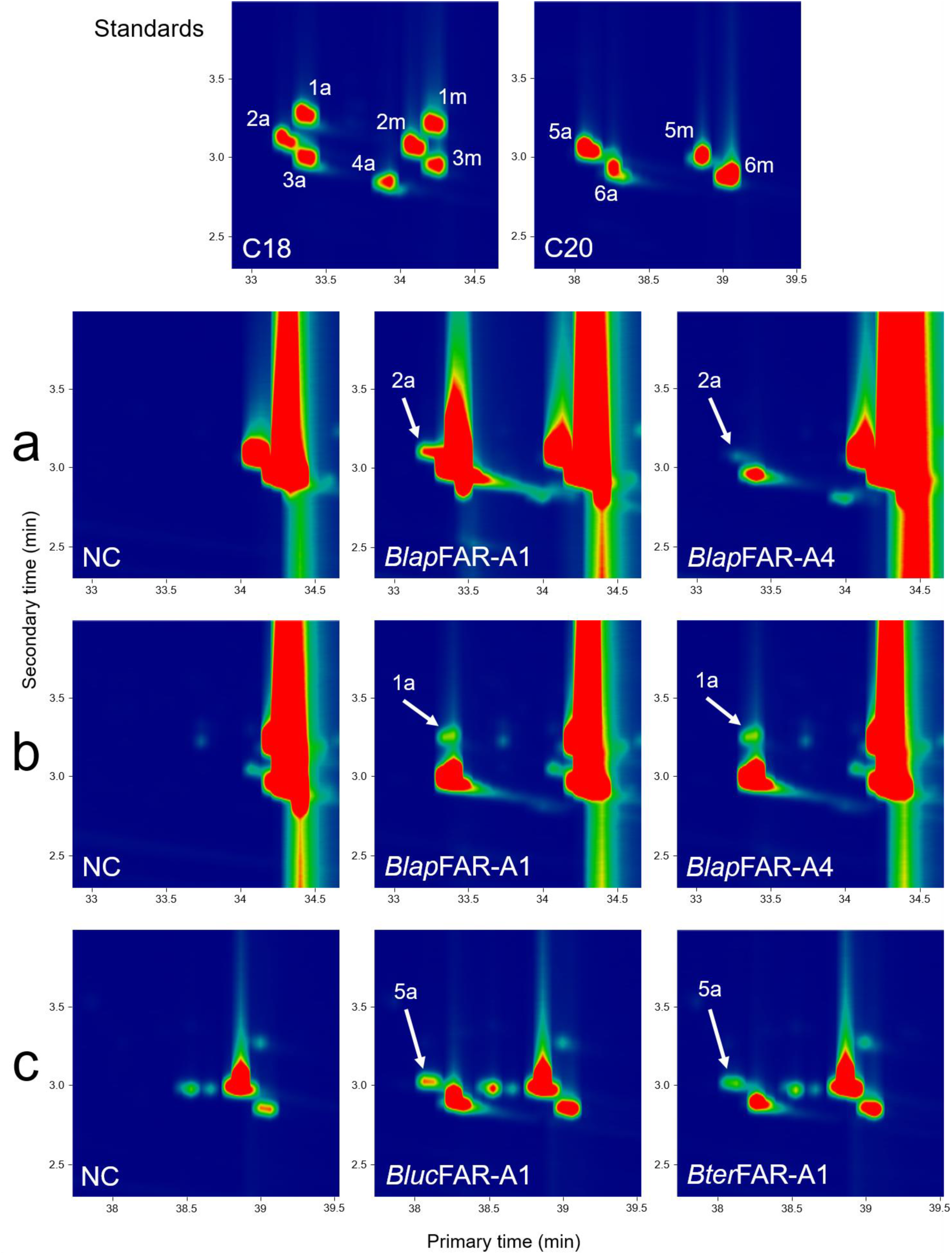
GC×GC-MS chromatograms of extracts from yeast strains supplemented with *Z*9,*Z*12-18:2 (**a**), *Z*9,*Z*12,*Z*15-18:3 (**b**) and *Z*15-20:1 (**c**) acyls. NC, negative control (yeast carrying empty vector); 1a, *Z*9,*Z*12,*Z*15-18:3OH; 1m, *Z*9,*Z*12,*Z*15-18:3Me; 2a, *Z*9,*Z*12-18:2OH; 2m, *Z*9,*Z*12-18:2Me; 3a, *Z*9-18:1OH; 3m, *Z*9-18:1Me; 4a, 18:OH; 5a, *Z*15-20:1OH; 5m, *Z*15-20:1Me; 6a, 20:OH; 6m, 20:Me. Selected masses: (**a**) 55+67, (**b**) 55+79, (**c**) 55+74, standards 55+67+74+79.

**Supplementary Figure 11.**
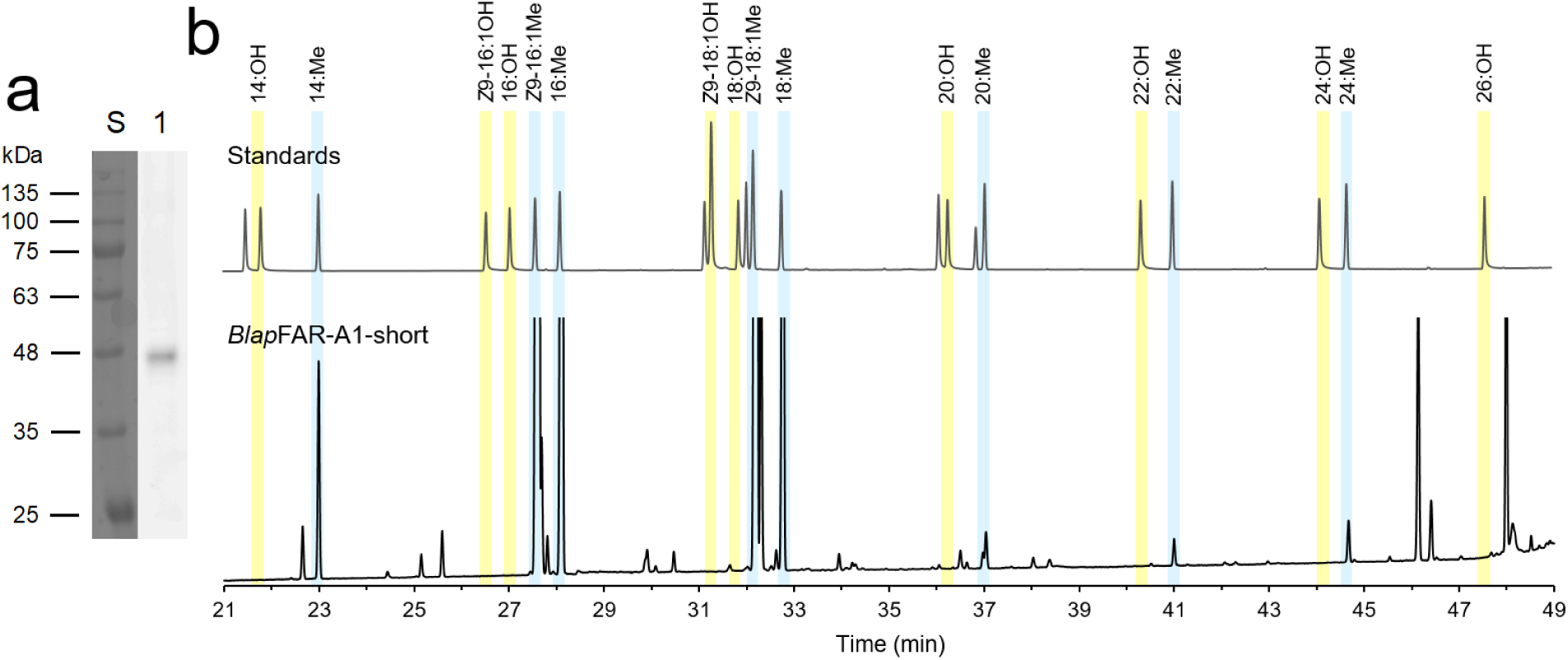
(**a**) Western blot analysis of cell lysate of yeast transformed with *Blap*FAR-A1-short. *Lanes*: S, protein standard (VI, AppliChem); 1, yeast carrying plasmid with the sequence of *Blap*FAR-A1-short. (**b**) GC-FID chromatogram of an extract from the yeast strain expressing *Blap*FAR-A1-short.

**Supplementary Figure 12.**
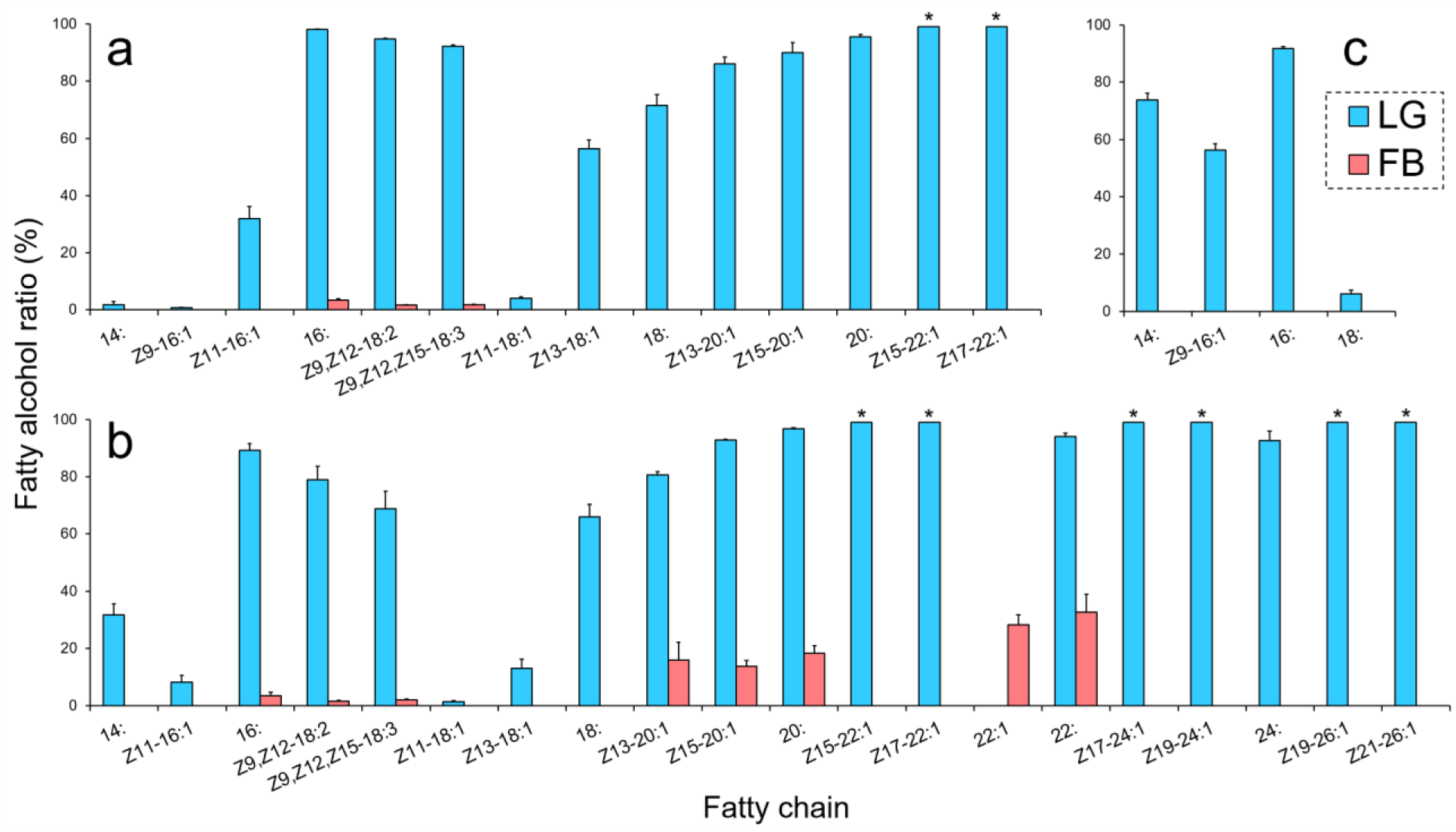
Fatty alcohol ratios in LGs and FBs of 3-day-old *B. lucorum* (**a**), *B. terrestris* (**b**) and *B. lapidarius* (**c**) males. Ratios >99% are marked with asterisks. In the case of 22:1 in FB of *B. terrestris*, the assignment of individual alcohol isomers was not possible due to low amount. See Methods for a description of fatty alcohol ratio calculation.

**Supplementary Table 1.** Predicted protein sequence lengths and conserved domains detected in predicted FAR coding regions via Conserved Domain Database search. The presence of a domain or conserved feature is marked with “+”.

**Supplementary Table 2.** Genomic linkage group or scaffold and orthology group of *B. terrestris* and *A. mellifera* FARs. *B. terrestris* FARs functionally characterized in this study and *A. mellifera* FAR previously functionally characterized are highlighted in green; FAR-A gene orthologs are highlighted in orange; genomic scaffolds not placed into linkage groups are grey.

| *Bombus terrestris* |  |  |
| --- | --- | --- |
| NCBI accession number | orthologous group | linkage group/scaffold |
| XP_020721700.1 | FAR-A | LG B12 |
| XP_020723892.1 | FAR-A | GroupUn1136 |
| XP_020723935.1 | FAR-A | GroupUn1199 |
| XP_003403375.2 | FAR-A | GroupUn1674 |
| XP_003402269.1 | FAR-A | GroupUn17 |
| XP_012175487.2 | FAR-A | GroupUn3510 |
| XP_012175505.2 | FAR-A | GroupUn4066 |
| XP_020723961.1 | FAR-A | GroupUn4067 |
| XP_012175509.2 | FAR-A | GroupUn4179 |
| XP_020723448.1 | FAR-A | GroupUn442 |
| XP_020723970.1 | FAR-A | GroupUn4704 |
| XP_003403427.2 | FAR-A | GroupUn4997 |
| XP_020723325.1 | FAR-A | GroupUn51 |
| XP_012174003.2 | FAR-A | GroupUn56 |
| XP_020723511.1 | FAR-A | GroupUn610 |
| XP_003402802.1 | FAR-A | GroupUn648 |
| XP_012174533.1 | FAR-A | GroupUn679 |
| XP_020723568.1 | FAR-A | GroupUn679 |
| XP_003403231.1 | FAR-A | GroupUn989 |
| XP_012175211.1 | FAR-A | GroupUn989 |
| XP_012175212.1 | FAR-A | GroupUn989 |
| XP_012175213.2 | FAR-A | GroupUn989 |
| XP_020723847.1 | FAR-A | GroupUn989 |
| XP_020723849.1 | FAR-A | GroupUn989 |
| XP_003394733.1 | FAR-A | LG B04 |
| XP_020720557.1 | FAR-B | LG B01 |
| XP_012174769.1 | FAR-C | GroupUn732 |
| XP_003402940.2 | FAR-D | GroupUn732 |
| XP_003399880.1 | FAR-E | LG B12 |
| XP_003399881.1 | FAR-F | LG B12 |
| XP_020721716.1 | FAR-G | LG B12 |
| XP_020721715.1 | FAR-H | LG B12 |
| XP_003399944.1 | FAR-I | LG B12 |
| XP_020720279.1 | FAR-J | LG B01 |
| XP_012173522.1 | FAR-K | LG B17 |
| *Apis mellifera* |  |  |
| NCBI accession number | orthologous group | linkage group/scaffold |
| XP_006565270.2 | FAR-A | LG12 |
| NP_001314876.1 | FAR-B | LG1 |
| XP_006570570.1 | FAR-C | LG6 |
| XP_006570601.2 | FAR-D | LG6 |
| ADI87409.1 | FAR-E | LG12 |
| ADI87411.1 | FAR-F | LG12 |
| NP_001180219.1 | FAR-G | LG12 |
| NP_001229455.1 | FAR-H | LG12 |
| XP_006566994.2 | FAR-I | LG12 |
| NP_001177849.1 | FAR-J | Un |
| XP_001120449.2 | FAR-K | LG5 |

**Supplementary Table 3.**
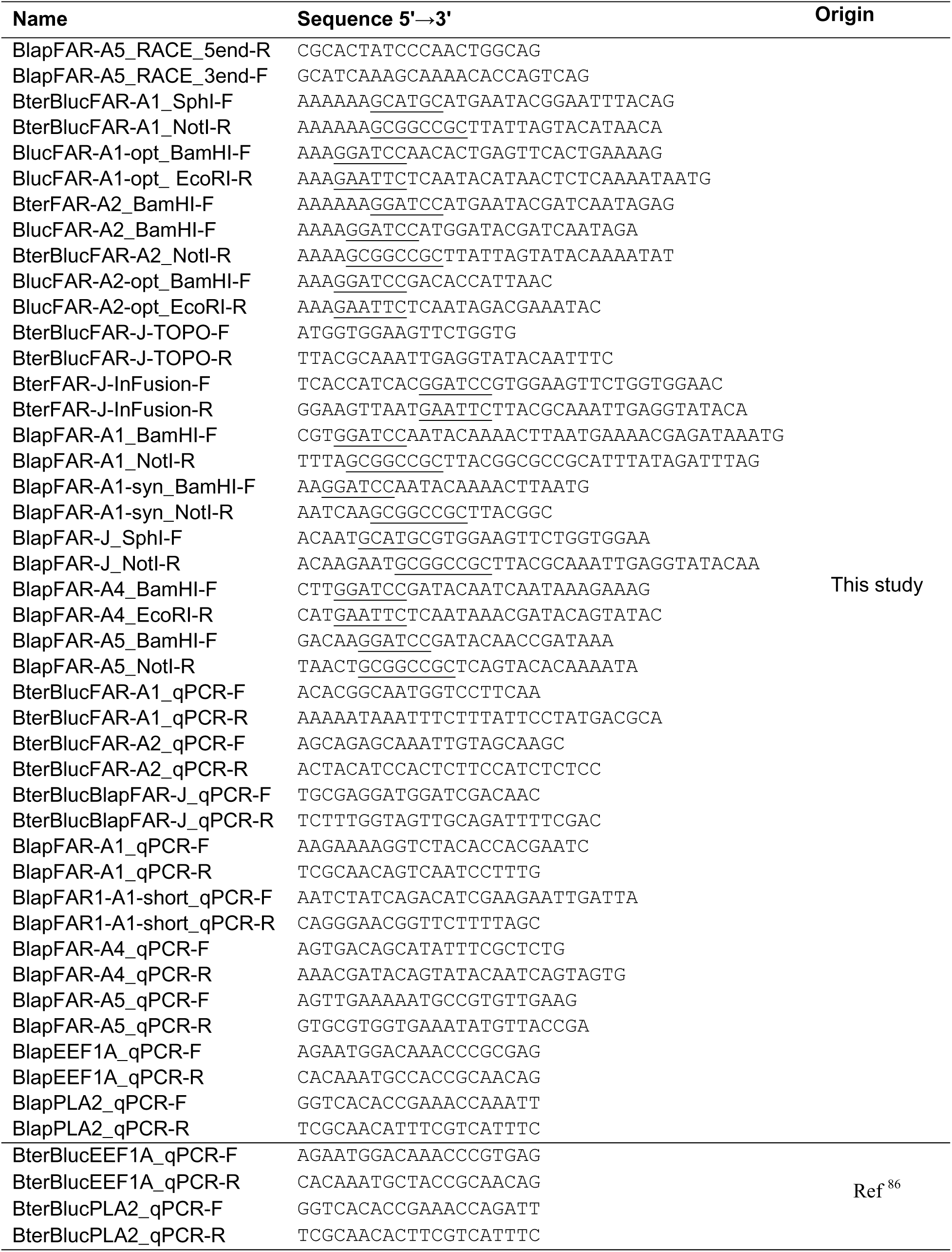
Primers used in this study. Restriction sites are underlined.

**Supplementary Table 4.** Protein sequence identities of bumblebee male marking pheromone (MMP)-biosynthetic FAR candidates. The colors indicate % identity (red: highest, blue: lowest).

|  | 1 | 2 | 3 | 4 | 5 | 6 | 7 | 8 | 9 | 10 |
| --- | --- | --- | --- | --- | --- | --- | --- | --- | --- | --- |
| 1: BterFAR-J_XP_020720279.1 | 100 |  |  |  |  |  |  |  |  |  |
| 2: BlucFAR-J_contig26150_9365 | 99.65 | 100 |  |  |  |  |  |  |  |  |
| 3: BlapFAR-J_contig12250 | 97.52 | 97.23 | 100 |  |  |  |  |  |  |  |
| 4: BlapFAR-A5_contig13763_4377 | 32.8 | 32.8 | 32.8 | 100 |  |  |  |  |  |  |
| 5: BlapFAR-A4_contig43815... | 34 | 34 | 34 | 78.77 | 100 |  |  |  |  |  |
| 6: BterFAR-A2_XP_003402269.1 | 32.41 | 32.41 | 32.41 | 77.78 | 79.37 | 100 |  |  |  |  |
| 7: BlucFAR-A2_contig55206_29748 | 32.8 | 32.8 | 32.8 | 75.79 | 77.98 | 94.84 | 100 |  |  |  |
| 8: BlapFAR-A1_contig44207 | 33.9 | 33.9 | 34.11 | 57.75 | 56.69 | 54.56 | 54.99 | 100 |  |  |
| 9: BterFAR-A1_XP_003394733.1 | 33.6 | 33.6 | 33.6 | 56.65 | 57.26 | 53.83 | 52.82 | 60.9 | 100 |  |
| 10: BlucFAR-A1_contig47309 | 33.4 | 33.4 | 33.4 | 56.85 | 57.26 | 53.83 | 52.82 | 61.11 | 99.4 | 100 |

**Supplementary Table 5.** Comprehensive GC analysis of fatty alcohols and transesterifiable fatty acyls in LGs of 3-day-old *B. terrestris, B. lucorum* and *B. lapidarius* males. The acyls were determined as corresponding methyl esters. *nd*, not detected.

| Tissue |  | Labial gland |  |  |  |  |
| --- | --- | --- | --- | --- | --- | --- |
| Species | *B. lapidarius* |  | *B. lucorum* |  | *B. terrestris* |  |
| Fatty chain | Amount (µg per tissue) |  |  |  |  |  |
|  | Acyl | Alcohol | Acyl | Alcohol | Acyl | Alcohol |
| Z7-12: | *nd* | *nd* | 0.84 ± 0.07 | *nd* | *nd* | *nd* |
| Z9-12: | *nd* | *nd* | *nd* | *nd* | 1.58 ± 0.08 | *nd* |
| 12: | 0.61 ± 0.04 | *nd* | 122 ± 4 | *nd* | 406 ± 24 | *nd* |
| Z5-14:1 | *nd* | *nd* | 2.39 ± 0.14 | *nd* | *nd* | *nd* |
| Z7-14:1 | 0.19 ± 0.02 | *nd* | *nd* | *nd* | *nd* | *nd* |
| Z9-14:1 | 2.13 ± 0.57 | *nd* | 1419 ± 192 | *nd* | 103 ± 18 | *nd* |
| 14: | 1.87 ± 0.21 | 4.70 ± 1.00 | 28.7 ± 2.9 | 0.45 ± 0.28 | 47.5 ± 4.7 | 19.5 ± 2.3 |
| Z7-16:1 | *nd* | *nd* | 14.2 ± 2.1 | *nd* | 5.59 ± 0.30 | *nd* |
| Z9-16:1 | 2236 ± 447 | 2577 ± 416 | 72.2 ± 5.7 | 0.44 ± 0.09 | 32.6 ± 0.6 | *nd* |
| Z11-16:1 | 15.8 ± 2.0 | *nd* | 3.40 ± 0.90 | 1.47 ± 0.56 | 37.7 ± 4.0 | 2.94 ± 0.63 |
| 16: | 157 ± 28 | 1560 ± 216 | 6.53 ± 0.47 | 313 ± 52 | 34.8 ± 1.9 | 263 ± 46 |
| Z9,Z12-18:2 | 10.6 ± 2.5 | *nd* | 2.50 ± 0.53 | 41.3 ± 8.2 | 13.9 ± 1.2 | 48.1 ± 10.0 |
| Z9,Z12,Z15-18:3 | 38.7 ± 6.8 | *nd* | 18.7 ± 4.6 | 201 ± 42 | 57.7 ± 0.5 | 119 ± 31 |
| Z9-18:1 | 119 ± 29 | *nd* | 57.0 ± 9.2 | *nd* | 129 ± 7 | *nd* |
| Z11-18:1 | 50.4 ± 10.7 | *nd* | 6.44 ± 0.87 | 0.25 ± 0.07 | 14.0 ± 1.3 | 0.18 ± 0.05 |
| Z13-18:1 | *nd* | *nd* | 3.85 ± 0.80 | 4.48 ± 0.45 | 47.5 ± 3.3 | 6.50 ± 1.56 |
| 18: | 7.27 ± 1.51 | 0.41 ± 0.02 | 5.16 ± 0.29 | 11.9 ± 2.4 | 8.35 ± 0.98 | 14.7 ± 1.4 |
| Z11-20:1 | *nd* | *nd* | 0.70 ± 0.03 | *nd* | 2.81 ± 0.51 | *nd* |
| Z13-20:1 | *nd* | *nd* | 1.02 ± 0.20 | 5.79 ± 0.76 | 2.65 ± 0.40 | 10.1 ± 1.4 |
| Z15-20:1 | *nd* | *nd* | 1.01 ± 0.23 | 8.77 ± 2.37 | 15.5 ± 1.3 | 182 ± 23 |
| 20: | 0.21 ± 0.04 | *nd* | 0.60 ± 0.05 | 11.9 ± 1.4 | 1.31 ± 0.08 | 35.9 ± 6.9 |
| Z15-22:1 | *nd* | *nd* | *nd* | 11.8 ± 2.8 | *nd* | 88.1 ± 9.5 |
| Z17-22:1 | *nd* | *nd* | *nd* | 1.88 ± 0.24 | *nd* | 14.9 ± 1.4 |
| 22: | *nd* | *nd* | 0.21 ± 0.09 | *nd* | 0.81 ± 0.03 | 12.2 ± 2.6 |
| Z17-24:1 | *nd* | *nd* | *nd* | *nd* | *nd* | 4.08 ± 0.37 |
| Z19-24:1 | *nd* | *nd* | *nd* | *nd* | *nd* | 6.46 ± 1.42 |
| 24: | *nd* | *nd* | *nd* | *nd* | 0.38 ± 0.10 | 4.79 ± 1.14 |
| Z19-26:1 | *nd* | *nd* | *nd* | *nd* | *nd* | 5.44 ± 1.17 |
| Z21-26:1 | *nd* | *nd* | *nd* | *nd* | *nd* | 7.51 ± 2.29 |
| 26: | *nd* | *nd* | *nd* | *nd* | 0.90 ± 0.30 | *nd* |
| 28: | *nd* | *nd* | *nd* | *nd* | 0.12 ± 0.03 | *nd* |

**Supplementary Table 6.** Comprehensive GC analysis of fatty alcohols and transesterifiable fatty acyls in FBs of 3-day-old *B. terrestris, B. lucorum* and *B. lapidarius* males. The acyls were determined as corresponding methyl esters. *nd*, not detected.

| Tissue |  | Fat body |  |  |  |  |
| --- | --- | --- | --- | --- | --- | --- |
| Species | *B. lapidarius* |  | *B. lucorum* |  | *B. terrestris* |  |
| Fatty chain | Amount (µg per tissue) |  |  |  |  |  |
|  | Acyl | Alcohol | Acyl | Alcohol | Acyl | Alcohol |
| 12: | 76.3 ± 17.0 | nd | 36.3 ± 10.6 | nd | 120 ± 34 | nd |
| Z5-14:1 | nd | nd | 2.60 ± 0.58 | nd | 0.84 ± 0.21 | nd |
| Z7-14:1 | 5.02 ± 0.92 | nd | 1.24 ± 0.36 | nd | 0.60 ± 0.22 | nd |
| Z9-14:1 | 20.4 ± 5.2 | nd | 58.1 ± 16.1 | nd | 106 ± 21 | nd |
| 14: | 128 ± 30 | nd | 158 ± 45 | nd | 456 ± 105 | nd |
| Z7-16:1 | nd | nd | 25.6 ± 7.3 | nd | 27.4 ± 5.7 | nd |
| Z9-16:1 | 817 ± 60 | nd | 67.5 ± 16.3 | nd | 134 ± 40 | nd |
| Z11-16:1 | 20.2 ± 2.7 | nd | 11.9 ± 6.5 | nd | 136 ± 22 | nd |
| 16: | 662 ± 112 | nd | 430 ± 108 | 13.82 ± 4.39 | 1333 ± 194 | 42.7 ± 11.8 |
| Z9,Z12-18:2 | 74.0 ± 7.5 | nd | 76.3 ± 18.2 | 1.12 ± 0.30 | 303 ± 37 | 4.35 ± 1.33 |
| Z9,Z12,Z15-18:3 | 245 ± 49 | nd | 597 ± 132 | 9.81 ± 2.32 | 1678 ± 435 | 32.2 ± 10.7 |
| Z9-18:1 | 788 ± 111 | nd | 1217 ± 269 | nd | 2951 ± 487 | nd |
| Z11-18:1 | 134 ± 8 | nd | 40.4 ± 19.3 | nd | 232 ± 30 | nd |
| Z13-18:1 | nd | nd | 31.7 ± 15.7 | nd | 69.5 ± 15.5 | nd |
| 18: | 60.8 ± 8.8 | nd | 119 ± 19 | nd | 224 ± 34 | nd |
| Z11-20:1 | nd | nd | 1.33 ± 0.39 | nd | 5.29 ± 0.68 | nd |
| Z13-20:1 | nd | nd | 0.66 ± 0.34 | nd | 4.40 ± 1.59 | 0.79 ± 0.47 |
| Z15-20:1 | nd | nd | 0.18 ± 0.12 | nd | 4.01 ± 1.56 | 0.56 ± 0.11 |
| 20: | 5.02 ± 0.21 | nd | 3.37 ± 1.06 | nd | 9.97 ± 2.24 | 2.10 ± 0.79 |
| Z13-22:1 | nd | nd | nd | nd | 0.54 ± 0.19 | nd |
| Z15-22:1 | nd | nd | nd | nd | 1.50 ± 0.49 | nd |
| Z17-22:1 | nd | nd | nd | nd | 0.15 ± 0.07 | nd |
| 22:1 (sum) | nd | nd | nd | nd |  | 0.81 ± 0.35 |
| 22: | nd | nd | 0.17 ± 0.08 | nd | 1.64 ± 0.56 | 0.78 ± 0.46 |
| 24: | nd | nd | nd | nd | 0.65 ± 0.35 | nd |
| 26: | nd | nd | nd | nd | 0.92 ± 0.50 | nd |
| 28: | nd | nd | nd | nd | 1.07 ± 0.64 | nd |

**Supplementary Data 1:** Synthetic nucleotide sequences. Restriction sites are underlined.

*Bluc*FAR-A1-opt (codon-optimized for yeast)

<u>GGATCC</u>AACACTGAGTTCACTGAAAAGTCTAACAAGGTCAACTCCATCGAAGGTTTCTACGCTGGTACAGGTATCTTTATCACAGGTGCCTCAGGTTTTGTCGGTAAAGGTTTGTTGGAAAAGTTGATCAGAGTTTGTCCTAGAATCGTTGTATTATTCATCTTGGTTAGACCAAAGAAACATCAAACAATGGAACAAAGATACAAGGAAATCATGGATGACCCTATCTTTGATGACATCAAAGCTAAGAATCCATCCGCATTGAAAAAGGTCCATCCTGTTGAAGGTGACATTTCCTTACCAAAGTTGGGTTTGAGTCAAGAAGATAGAAACATGTTGATAGAAAACGTCAACATCTTGTTTCACGTTGCTGCATCTTTGAACTTCAAGGAACCATTGAACGCCGCTGTAAATACTAACGTCAAGGGTACATTTTCTATAATCGAATTGTGTAACGAATTGAAGCATGTTATATCAGCTGTACACGTCTCTACAGCATATTCAAATGCCAACTTGCCTGAAATAGAAGAAAAGGTTTACTCCACTATCTTACAACCATCTTCAGTAATTGAAACATGCGACAGTTTGGATAAGGAATTGATTAAGTTGTTGGAAGAAAGAATTTTGAAAATACATCCTAACACGTACACCTTCACGAAGAATTTGGCAGAACAAATCTTGTCCAGTTCTTCAACTAACTTCCCAATAGCAATCGTTAGACCTTCTATCATTTCCGCCAGTTTAAAAGAACCATGTCCTGGTTGGTTGGGTAATATTACAGCCCACATAGCTTTGGGTTTGTTTATTTCAAGAGGTTTCGCCAAGATCACCTTAGCTAACCCTGACACTATCACAGATACCGTACCATTAGACTATGTCGTTGATACAATTTTGTGTGCAGCCTGGCATGTTACCTTGCACAGAGATATGAACGTTAAGGTATACAACTGCACCAATAACGCCAGACCAATTAATTACGGTGAATTGAAGGACACTTTTGTCAAGTACGCTATTCAAATACCTATGGATGGTTTAGTTTGGTATCCATGTTGCGCAATGGTTTCTAACAGATACGTATACTCAATCTTGACCTTATTCTTGCATACTTTGCCTGCTTTTATTATGGATATTTTCTTAAGATTGCAAGGTTCAAAGCCAAGAATGATGAAGATCTCTAAGTACTACGATACAATGTCAATCGTTACCAACTACTTCTCCACTAGACAATGGAGTTTCAAAAAGGATAACGTTATTAATATGATGAAAGAAGTCAAGACTTTGGAAGATTCTGACATCGTTAGATTAGATTTGCAAGATATGGACTGGGATAAGTACATCGCTATATGCGTTATCGGTATCAAAAAGTTTATTTTCAAAGAAGACCCAAAGTCCTTAGATGCTGCATTGAGAAGATTGAGTATCTTTTACTGGATTCATCAAATGACTAAAGCCTTCGCTATTATTATCTTATTGACCATTATTTTGAGAGTTATGTATTGA<u>GAATTC</u>

*Bluc*FAR-A2-opt (codon-optimized for yeast)

<u>GGATCC</u>GACACCATTAACAGAGAAAAGAACGAAAACGCCATTAACAAGGGTTTGAACAAGTTGAATACATTAGAAGAATTTTACGTCGGTAGTGGTATTTTGTTAACTGGTGCAACAGGTTTTGTTGGTAAAGCTGTTTTGGAAAAGTTGATCAGAATGTGTCCAAGAATTGCTGCAATTTTCTTGTTGTTTAGACCAAAGACTGATGAAACAATCGAACAAAGATTCAAGAAATTGATTGATGATCCAATCTATGATGATATCAAGGCAAAGCATCCATCAACTTTGTCTAGAGTTTATCCAATGAGAGGTGACTTGTCATTGCCAGATTTGGGTTTGTCTAGAGAAGATAGAAATTTGTTGTTGGAAAAGGTTAACATCGTTTTCCATGCTGCAGCTACTGTTATGTTTAATGAACCATTGCAAGTTACAATTAATGTTAACACTAAAGGTACAGCTAGAGTTATTGATTTGTGGAACGAATTGAAGCATCCAATCTCATTCGTTCATGTTTCAACTGCATTTTCTAACGCTAACATCCATGAAATCGGTGAAAGAGTTTACACTACATCATTGAAACCATCTGAAGTTATTGATATTTGTAATAAGTTTGATAAAACATCTATTAATCAAATCGAAAAGAAAATTTTGAAAACTTATCCAAATATCTATACTTTTTCTAAAAATTTGGCAGAACAAATCGTTGCTTCTAACTGTAAGGATATGCCAGTTGCTATTGTTAGACCATCAGTTATTGGTCCATCTATGGAAGAACCATGTCCAGGTTGGATTCAAAACATCTCTGCAATCACTGGTATCATGGTTTTGATCGGTAGAGGTTGTGCTACAGGTATTAGAGGTAGAAGAGATGGTAGAGTTGATGTTGTTCCATTGGATTACGTTGTTGATATGATCATCTGTACTGCATGGCATGTTACATTGCATCCAAAGCATGAAGTTAAGGTTTACAACTGTACATCTTCAGCTAACCCAATTAGATGGGGTCAAATGCAACAATTGGTTTTGAAGCATTCAAGAGAAACTCCATTAAACGATACATTGTGGTATCCAAGATGTCCAATCATCGCAAATAAGTACATTTTCAATGTTTTATGTGTTATCCCATACGTTTTGCCAGCTTTTATTATCGATATTTTCTTGAGATTGAGAGGTTCTAAGCCAATCATGATGAAGTTGTTGAAGTTCGGTTACAAGTTGTCTACTTCAGTTTCTCATTTCACTATGAACGAATGGACATTCCAAAGAGATAACTGTTCAGATTTGGCATCTAAGGTTAAGATGTTGCATGATTCAGATATGGTTAAATTAGATTTGAGAGATATGAAGTGGGAAAAGTACATCGTTATATATTTGATGGGTATTAGAAAGTTTATTTTGAAACAAGAATTTCAACCAACAGCTAGACAAAGATTGTCTAGATTGTACTGGATTCATCAAATCACTAAGATTTCAGGTATCATCAGTTTGTTATGGATTATTTTGTATTTCGTCTATTGA<u>GAATTC</u>

*Blap*FAR-A1

<u>GGATCC</u>AATACAAAACTTAATGAAAACGAGATAAATGAAAAATTACGTAATGTGAATTCCATTGGGGGATTCTACGCCGGAACTGGAATTCTTATTACTGGTGGGACAGGTTTCGTGGGCAAAGGACTCCTCGAAAAACTGATACGCACGTGTTCACACATCGCTGCTATTTTTATATTGATCCGTCCGAAACGTAACCAAACGATAGAACAACGATTTAAGAAGATAATAGATGATCCGATTTTCGATGGTGTCAGAGCACAAAACCCAGCAATTTTCTATAAAATTCATCTCGTGGAGGGCGACGTGACTCTACCAGATTTAGGTCTTTTGCAAAAAGACAGAGATATGTTGATAGAGAATGTAAACATAGTGTTCCACATTGCGGCCACTATAAATTTCCATCAACCATTGGATATGATTGTCAATGTAAATGTGAAAGGTACCGCTAATATTATCAAACTGTGCAAGGAACTCAAGCATGTAATTAGCGTTGTCTATGTGAGCACAGCTTACAGTAATCCGAATCTATCAGACATCGAAGAAAAGGTCTACACCACGAATCTAGATCCCTCTCTCGTGATGGATATATGCGACCGACAAGACAAAGAATTGATTAATCTGATCGAAGAAAGAATTTTAAAAACGCATCCGAACACATACACGTTCACCAAGAATCTTGCAGAGCAGACAATATCCAACAATAGCAAAGGATTGACTGTTGCGATAGTGCGACCAAGTATAATTTCTTCCTCGCTAAAAGAACCGTTCCCTGGTTGGTTGGTATCTTTTGCTGGACAATCAGGTATCTTCAAGAATATCGGCAATGGTATGGCAAAAGTACTATTGGGTAGGGGAGATGTAATATCAGATATAGTGCCTGTTGATTATGTAGTCGACGCGATAATGTGTGCCGCGTGGCACGTCACGCTACAAATTGATAATAATGTCAAAGTTTACAACTGTACGAGCAGCGCACGTCCCATCAAATTGGGTGAAATCGTAAATATCTTCGTGGAATGTAGCAGAGAAATACCGATGAAAAATACGTTGTGGTATCCGAGTTGTACGATAGTAGCAAACAGATTTGTTTACAATGTACTGAATATACTTCTAAATGTTTTACCTGCGTTTGCCGTGGATATCTTTTTAAGGCTTCGAGGTGGTAAACCAATGGCAATGAATATGAACAAATATTACAATAAATTGGTCGTAGCGACAAGCTACTTCAACTCGAATGAATGGTCCTTCAAAAGAGATAACATTGCCGATATGATAAACAAGGTGAATACCTTGGAAGATGGAAATATTGTTAAACTGGACTTGCAGGATATGGTTTGGAGGAAATATATAGCAAATTACTTGGCGGGAATTAAGAAATTTATTCTGAAAGAAGACCCTAAATCTATAAATGCGGCGCCGTAA<u>GCGGCCGC</u>

